# Insights about variation in meiosis from 31,228 human sperm genomes

**DOI:** 10.1101/625202

**Authors:** Avery Davis Bell, Curtis J. Mello, James Nemesh, Sara A. Brumbaugh, Alec Wysoker, Steven A. McCarroll

## Abstract

Meiosis, while critical for reproduction, is also highly variable and error prone: crossover rates vary among humans and individual gametes, and chromosome nondisjunction leads to aneuploidy, a leading cause of miscarriage. To study variation in meiotic outcomes within and across individuals, we developed a way to sequence many individual sperm genomes at once. We used this method to sequence the genomes of 31,228 gametes from 20 sperm donors, identifying 813,122 crossovers, 787 aneuploid chromosomes, and unexpected genomic anomalies. Different sperm donors varied four-fold in the frequency of aneuploid sperm, and aneuploid chromosomes gained in meiosis I had 36% fewer crossovers than corresponding non-aneuploid chromosomes. Diverse recombination phenotypes were surprisingly coordinated: donors with high average crossover rates also made a larger fraction of their crossovers in centromere-proximal regions and placed their crossovers closer together. These same relationships were also evident in the variation among individual gametes from the same donor: sperm with more crossovers tended to have made crossovers closer together and in centromere-proximal regions. Variation in the physical compaction of chromosomes could help explain this coordination of meiotic variation across chromosomes, gametes, and individuals.

## Introduction

One way to learn about human meiosis has been to study how genomes are inherited across generations. DNA variation data are now available for millions of people and thousands of families; the locations of crossovers can be estimated from genomic segment sharing among relatives and linkage-disequilibrium patterns in populations^1–5^. Although these studies sample only the small number of reproductively successful gametes from individual humans, such analyses have revealed that average crossover number and crossover location vary among individual humans and associate with common variants (single nucleotide polymorphisms, SNPs) at many genomic loci^4, 6–10^.

Another powerful approach to studying meiosis is to directly visualize meiotic processes in individual cells. For example, technical innovations have made it possible to ascertain that homologous chromosomes in spermatocytes generally begin synapsis (their physical connection) near their telomeres^11–13^; to observe double-strand breaks (a subset of which progress to crossovers) by monitoring proteins that bind to such breaks^14–17^; and to detect adverse meiotic outcomes, such as chromosome mis-segregation^18–23^. Studies based on such methods have revealed substantial cell-to-cell variation, even among cells from the same individual, in features such as the physical compaction of meiotic chromosomes^24–26^.

More recently, human meiotic phenotypes have begun to be studied via genotyping or sequencing up to 100 gametes from one person, demonstrating that crossovers and aneuploidy can be ascertained from direct analysis of gamete genomes^27–31^. Despite these advances, it has not yet been possible to measure meiotic phenotypes genome-wide in many individual gametes from many people.

## Results

### A high-throughput single-sperm sequencing method

To this end, we developed a method called “Sperm-seq” with which the genomes of many individual sperm can be sequenced to low coverage quickly and simultaneously. To access the tightly compacted sperm genome, we decondense sperm nuclei using reagents that mimic the molecules with which the egg unpacks the sperm pronucleus (**Fig. 1a**, Methods). These decondensed sperm DNA “florets” are then encapsulated with barcoded beads in microfluidic droplets in which the sperm genomes are individually barcoded and amplified^32^. Each genomic sequence read has a barcode that reports its droplet—and thus gamete—of origin (**Fig. 1a**). We used this technique to sequence 31,228 sperm cells from 20 sperm donors (974-2,274 gametes per donor), sequencing a median of ∼1% of the haploid genome of each cell (Table 1); deeper sequencing allows coverage of ∼10% of a gamete’s genome.

**Figure 1.**
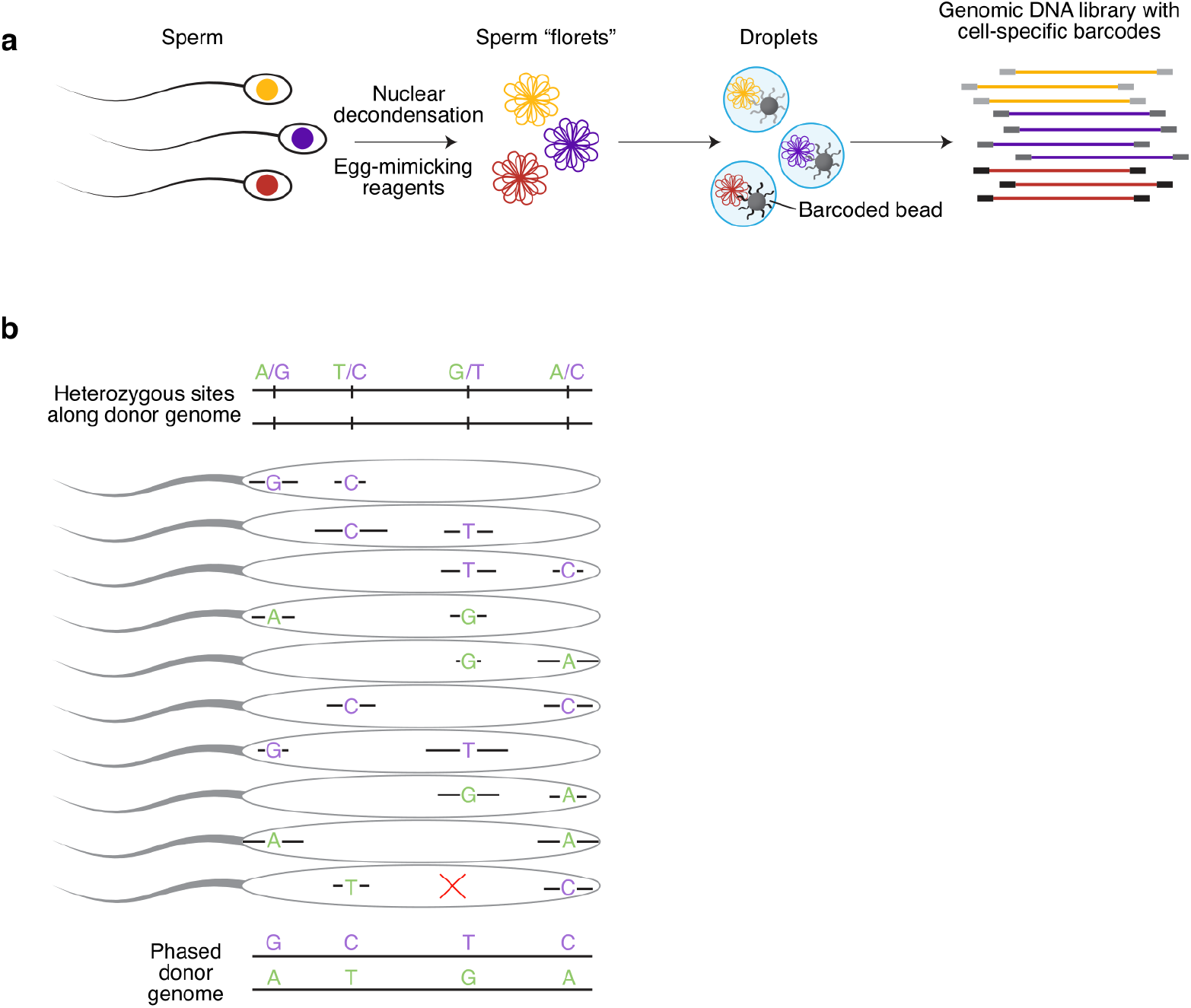
High-throughput single-sperm sequencing and chromosome-length haplotype phasing strategy. **a**, Sperm cells are decondensed into sperm DNA “florets,” encapsulated in droplets with barcoded beads^32^ and whole-genome amplified, followed by sequencing library preparation. **b**, Phasing strategy. Green and purple denote phase of allele (unknown before analysis). Each sperm cell carries one parental haplotype (green or purple) except where a recombination event separates consecutively observed SNPs (red “X” in bottom sperm). Because alleles from the same haplotype will tend to be observed in the same sperm cells, haplotypes are resolvable and can be assembled to whole-chromosome scale. Extended Data Fig. 1 evaluates phasing performance and illustrates use of phased haplotypes to identify cell barcodes associated with more than one sperm cell (cell doublets).

**Table 1.** Sperm donor and single-sperm sequencing characteristics and results.

| Donor | Ancestry* | Cells (number excluding cell and bead doublets) | Reads per cell (median, thousands) | Genome covered per cell (median, percent) | Heterozygous SNPs in genome (millions) | Unique heterozygous SNP alleles observed per cell (median, thousands) | Crossovers observed (total, thousands) | Crossovers per cell (mean) | Resolution of crossovers (kb, median) | Autosomal aneuploidy events (percent of cells)<sup>#</sup> | Sex chromosome aneuploidy events (percent of cells)<sup>#</sup> |
| --- | --- | --- | --- | --- | --- | --- | --- | --- | --- | --- | --- |
| **Overall** | -- | 31,228<sup>a</sup> | 211<sup>b</sup> | 1.0<sup>c</sup> | -- | 24.6<sup>b</sup> | 813<sup>a</sup> | 26.11<sup>c</sup> | 240<sup>d</sup> | 1.6<sup>b</sup> | 0.9<sup>b</sup> |
| NC1 | Eur. | 982 | 284 | 1.4 | 1.95 | 31.6 | 26 | 26.31 | 189 | 1.5 | 0.6 |
| NC2 | Eur. | 1,680 | 163 | 0.8 | 1.98 | 18.2 | 37 | 22.19 | 307 | 2.0 | 0.7 |
| NC3 | Eur. | 1,289 | 190 | 0.9 | 1.94 | 21.5 | 36 | 28.13 | 260 | 1.7 | 0.7 |
| NC4 | Eur. | 1,482 | 243 | 1.1 | 1.98 | 26.8 | 40 | 26.98 | 243 | 1.1 | 0.5 |
| NC6 | Afr. Am. | 1,370 | 154 | 0.8 | 2.53 | 23.8 | 38 | 27.57 | 253 | 0.7 | 0.3 |
| NC8 | As. | 1,663 | 304 | 1.5 | 1.81 | 30.9 | 45 | 26.98 | 229 | 3.1 | 0.5 |
| NC9 | As. | 1,894 | 245 | 1.2 | 1.79 | 25.6 | 53 | 27.98 | 231 | 0.8 | 1.5 |
| NC10 | As. | 1,154 | 224 | 1.1 | 1.82 | 23.3 | 29 | 24.99 | 257 | 1.5 | 0.3 |
| NC11 | Eur. | 1,930 | 202 | 1.0 | 1.92 | 22.8 | 50 | 25.82 | 242 | 1.3 | 0.4 |
| NC12 | Eur. | 2,145 | 179 | 0.9 | 1.91 | 20.6 | 51 | 23.76 | 270 | 1.2 | 1.7 |
| NC13 | Eur. | 1,514 | 259 | 1.2 | 1.92 | 28.3 | 41 | 27.19 | 202 | 0.9 | 1.0 |
| NC14 | Eur. | 1,336 | 296 | 1.4 | 1.92 | 32.4 | 36 | 26.65 | 175 | 2.5 | 1.2 |
| NC15 | Eur. | 1,702 | 211 | 1.0 | 1.93 | 23.2 | 42 | 24.80 | 268 | 1.0 | 0.9 |
| NC16 | Eur. | 1,785 | 241 | 1.2 | 1.92 | 26.9 | 42 | 23.78 | 227 | 2.2 | 1.3 |
| NC17 | Eur. | 1,504 | 220 | 1.0 | 1.94 | 23.8 | 39 | 25.92 | 250 | 2.1 | 0.7 |
| NC18 | Eur. | 1,589 | 170 | 0.8 | 1.93 | 18.4 | 42 | 26.48 | 317 | 1.2 | 0.6 |
| NC22 | Afr. Am. | 1,693 | 195 | 0.9 | 2.53 | 29.7 | 44 | 25.96 | 205 | 1.4 | 0.7 |
| NC25 | Afr. Am. | 2,274 | 175 | 0.8 | 2.47 | 25.8 | 62 | 27.31 | 211 | 2.8 | 1.8 |
| NC26 | Afr. Am., As. | 974 | 120 | 0.6 | 2.55 | 18.0 | 26 | 26.67 | 355 | 1.3 | 0.4 |
| NC27 | As. (?) | 1,268 | 267 | 1.3 | 1.96 | 29.2 | 34 | 26.80 | 199 | 1.7 | 0.6 |

\*As provided by sperm bank. Afr. Am., of African American ancestry; Eur., of European ancestry; As., of Asian ancestry; (?), conflicting ancestry information given.
<sup>#</sup>These numbers are the total number of aneuploidy events divided by the total number of cells multiplied by 100; cells can have more than one event.
<sup>a</sup> Sum across all cells from all sperm donors.
<sup>b</sup> Median or mean across all individual cells from all sperm donors (31,228 measurements summarized).
<sup>c</sup> Median or mean of aggregate metrics across samples (20 measurements summarized).
<sup>d</sup> Median across all crossovers (813,122 measurements summarized).

Data from so many individual gametes made it possible to infer individuals’ allelic haplotypes along the full length of every chromosome. We first identified the heterozygous sites in each donor’s genome using the Sperm-seq sequence reads (∼40x coverage per donor, Methods). Because each sperm chromosome is a mosaic of long segments derived from one or the other parental haplotype, the chromosomal phase of heterozygous sites could be inferred from the co-appearance patterns of alleles (of different SNPs) across many sperm cells (**Fig. 1b**, Methods). *In silico* simulations and comparisons to haplotypes from population-based analyses indicated that Sperm-seq assigned alleles to haplotypes with 97.5–99.9% accuracy (Extended Data Fig. 1a, Supplemental Text). These phased haplotypes made it straightforward to identify and remove from the analysis cell “doublets,” cases in which two sperm genomes were tagged with the same cell barcode, from the presence of both parental haplotypes at multiple loci across chromosomes (Extended Data Fig. 1b-d, Methods). We also identified surprising “bead doublets,” in which two beads’ barcodes appeared to have tagged the same gamete genome, as they reported identical genome-wide haplotypes (ascertained through different SNPs) (Extended Data Fig. 2a,b, Methods). Bead doublets were useful for evaluating the replicability of Sperm-seq data and analyses, which is usually impossible to do in inherently destructive single-cell molecular studies (Extended Data Fig. 2c-e).

### Recombination rate in sperm donors and sperm cells

Analysis of Sperm-seq data identifies crossover (recombination) events as transitions between parental haplotypes (**Fig. 2a**, Methods). We identified 813,122 crossovers in the 31,228 gamete genomes (mean 26.03 per gamete; 25,839-62,110 per sperm donor, Table 1). Crossover locations were inferred with a median resolution of 240 kb, and 9,746 (1.2%) were inferred at resolution finer than 10 kb (Table 1, Supplemental Text). In analysis of data from bead doublets, 95.6% of crossovers were detected in both cell barcodes; another 2.1% were near the ends of SNP coverage on chromosomes, where the power to detect crossovers is incomplete (Extended Data Fig. 2e). Estimates of crossover rate and location were robust to down-sampling to the same number of SNP observations in each cell (Extended Data Fig. 3, Methods).

**Figure 2.**
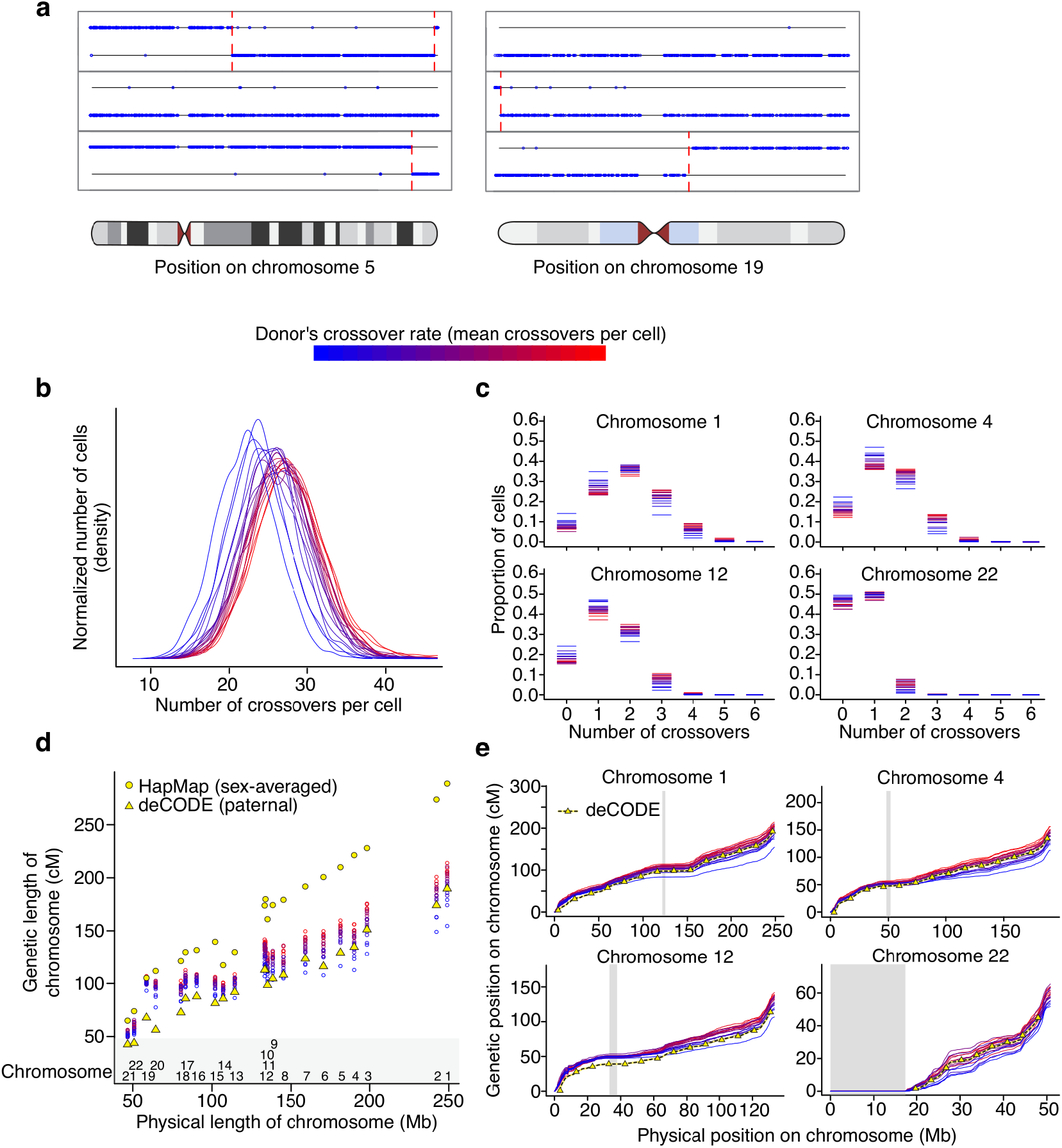
Crossover identification and recombination rate from single-sperm sequencing. **a**, Crossover calling strategy. Three example cells are shown for each of two chromosomes. After genome phasing, each allele observed at any heterozygous site (blue circle) in a cell is assigned to its haplotype of origin (horizontal lines); crossovers (vertical dashed red lines) are transitions between haplotypes. **b**, Density plot showing per-cell number of autosomal crossovers for all 31,228 cells and 813,122 total autosomal crossovers from 20 sperm donors (per-donor cell and crossover numbers as in Table 1, aneuploid chromosomes excluded from crossover analysis). Line colors, donor’s mean crossovers per cell from low (blue) to high (red). This same mean recombination rate-derived color scheme is used for donors in all figures. Recombination rate differs among donors (Kruskal–Wallis chi-squared = 3,665, *df* =19, *p* < 10^−300^). (Extended Data Fig. 5 shows this data cumulatively.) **c**, Per-chromosome crossover number in each of the 20 sperm donors (data as in (**b**) but segmented by chromosome). Extended Data Fig. 4 shows all 22 autosomes. **d**, Per-chromosome genetic map lengths from each donor, linkage-based genetic chromosome lengths from HapMap^5^ (includes crossover-rich female meioses), and pedigree-based chromosome lengths from deCODE^10^ (deCODE genetic maps stop 2.5 Mb from the ends of SNP coverage). **e**, Physical vs. genetic distances (for individualized sperm donor genetic maps and deCODE’s paternal genetic map) plotted at 500 kb intervals (hg38). Gray boxes denote centromeric regions (or centromeres and acrocentric arms). Extended Data Fig. 6 shows all 22 autosomes.

Crossovers, which create new allelic combinations, differ in genomic locations and average number among individual humans^2, 3, 6, 7, 9, 10^. The 20 individual sperm donors exhibited recombination rates ranging from 22.2 (95% confidence interval [CI] 22.0–22.4) to 28.1 (95% CI 27.9–28.4) crossovers per cell, consistent with earlier rate estimates from a few living children^2, 6–10^ or up to 100 sperm cells^29, 31^ (Table 1, **Fig. 2b,c**, Extended Data Figs. 4, 5). For each chromosome, the proportion of cells with each observed number of crossovers varied among individuals in the way predicted by their global crossover rate (**Fig. 2c**, Extended Data Fig. 4). The 813,122 inferred crossovers allowed us to generate genetic maps for each of the donors; these maps were broadly concordant with deCODE’s paternal genetic map previously estimated by genotyping thousands of families^10^ (**Fig. 2d,e;** Extended Data Fig. 6; Supplemental Text).

More variation was present at the single-cell level: the range in the routine number of crossovers per cell was 17 to 37 (1^st^ and 99^th^ percentiles, median across donors), with an across-cell standard deviation of 4.23 (median across donors). Crossover number could in principle be co-regulated nucleus-wide, as suggested by the correlation of crossover number across chromosomes observed in pedigrees^9^ and spermatocytes undergoing meiosis^25, 26^. In fact, individual gametes with fewer crossovers in half of their genome (the odd-numbered chromosomes) did tend to have fewer crossovers in the other half of their genome (Pearson’s *r* = 0.09, *p* = 8 × 10^−54^ with all gametes from all donors combined after within-donor normalization; Supplemental Text). (This point estimate greatly underestimates the true correlation of crossover number across chromosomes in spermatocytes, as any co-regulation of crossover number across chromosomes would occur in the spermatocyte, whose daughter gametes each have only a 50% chance of inheriting any given parental crossover.)

On any given chromosome, fewer cells had no crossovers or many crossovers than would be predicted by a model in which crossovers are independent, random events (Extended Data Fig. 7; Supplemental Text). This is consistent with biological constraint on crossover number, a major determinant of which is crossover interference (reviewed in ^33, 34^).

### Crossover location and interference

Crossovers are distributed non-uniformly along chromosomes, in patterns that vary at both fine scales (such as their recurrence in hotspots) and large scales (such as their concentration in sub-telomeric regions in male meiosis)^1, 4–7, 10, 35, 36^. Although the spatial resolution of most crossover inferences was not well suited for analyzing fine-scale selection of crossover sites (*e.g.,* hotspots), the large number of crossovers ascertained per sperm donor (25,839-62,110) made it possible to analyze variation in large-scale crossover placement.

Crossovers were concentrated in large regions of the genome (“crossover zones”) that were shared across donors (**Fig. 3**, Extended Data Figs. 8, 9). Zones in the sub-telomeric regions had the most crossovers, whereas regions close to the centromere had fewer crossovers, consistent with earlier findings^1, 4, 6, 10, 37^ (**Fig. 3a**). However, on the larger acrocentric chromosomes (chromosomes 13, 14, and 15), which do not perform crossovers in their *p* arms, each centromere-proximal zone had a crossover rate comparable to the most telomeric zone on the same chromosome.

**Figure 3.**
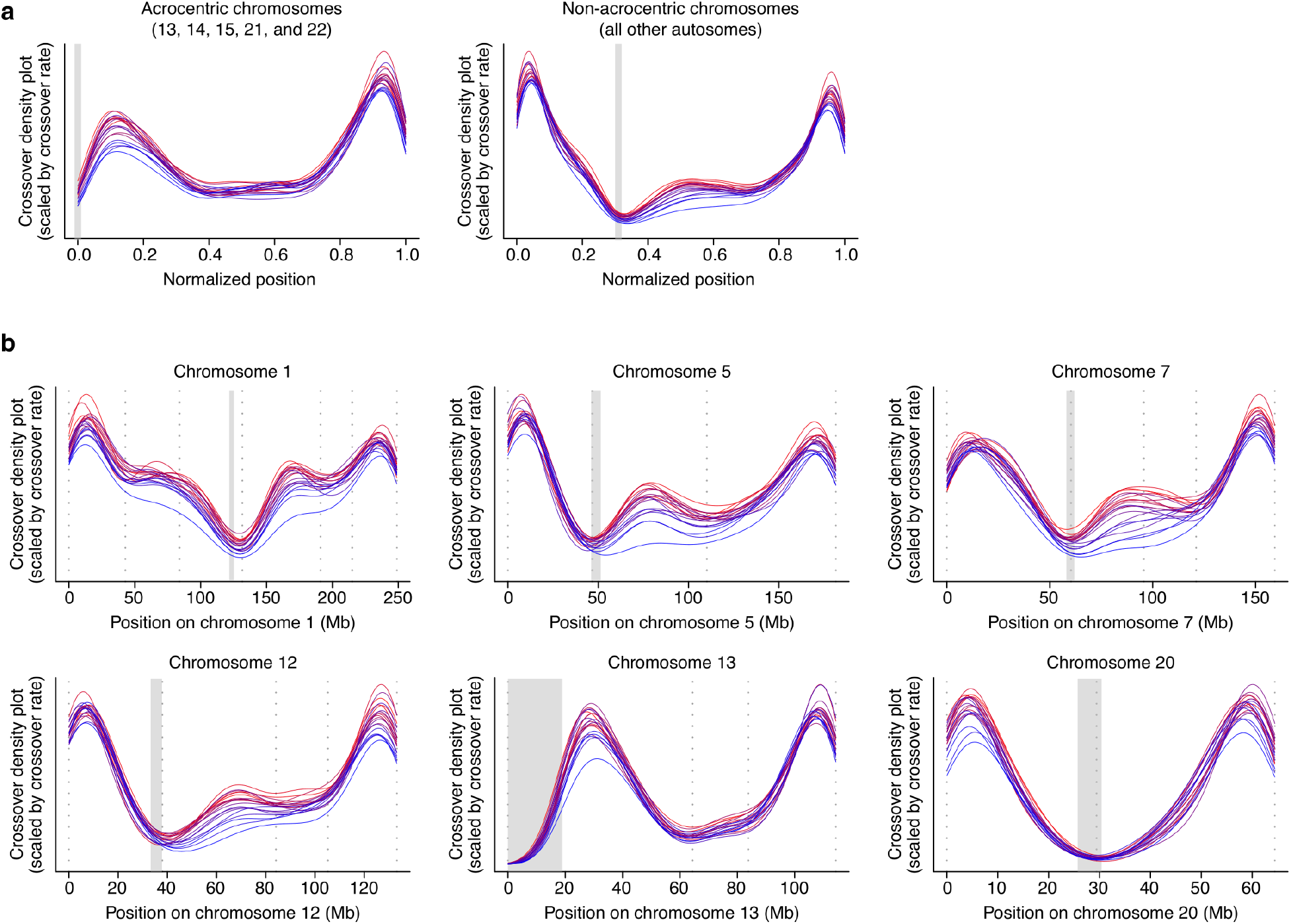
Crossover location patterns across chromosomes (in “crossover zones”). Each donor’s crossovers are plotted as a colored line; color indicates donor crossover rate as in Fig. 2.; gray boxes mark centromeres (or centromeres and acrocentric arms). The midpoint between the SNPs bounding the crossover was used as the single position for each crossover in all analyses. **a**, Crossover locations (density plot) on “meta-chromosomes.” All crossovers are plotted based on where they occurred in the chromosome arm. For acrocentric chromosomes, only the *q* arm was considered; for non-acrocentric chromosomes, the *p* and *q* arms were afforded space based on the proportion of the genome (in bp) they comprise. **b**, Each donor’s crossover location density plot for individual chromosomes (per-donor numbers in **Table 1**). The area under each curve is proportional to the crossover rate on that chromosome for each donor. Dotted gray vertical lines denote crossover zone boundaries (separating crossover-preferred regions, Extended Data Fig. 8). Extended Data Fig. 9 shows all 22 autosomes.

The crossover zones with the most variable usage (across people) were all adjacent to centromeres (**Fig. 3b**, Extended Data Fig. 9); individuals with high recombination rates used these zones much more frequently (crossover location patterns were robust to among-donor coverage differences, Extended Data Fig. 3c,e). Of the 10 crossover zones with crossover rates correlating most strongly with global recombination rate, all but one were centromere-proximal, and the exception was separated from the centromere by only one small zone. Consequently, the proportion of crossovers in the most distal zones of the chromosomes varied strikingly among individuals (Kruskal–Wallis chi-squared = 2,334, *df* = 19, *p* < 10^−300^) and was negatively correlated with recombination rate (Pearson’s *r* = −0.95, *p* = 2 × 10^−10^) (Extended Data Fig. 10a).

Crossover interference, which manifests in the tendency of crossovers to be further apart than expected by chance, occurs in humans^25, 31, 37–41^. The effect of crossover interference was visible in each of the 20 sperm donors: the distances between consecutive crossovers were greater in the observed data than when crossover locations were permuted across cells (Extended Data Figs. 11-15). The extent of crossover interference varied greatly among individual sperm donors (Kruskal–Wallis chi-squared = 4,316, *df* = 19, *p* < 10^−300^) and correlated inversely with a donor’s global recombination rate (Pearson’s *r* = −0.99, *p* = 9 × 10^−16^) (Extended Data Fig. 10b).

We estimated crossover placement and interference from the 180,738 chromosomes with exactly two crossovers to determine whether the relationships between these meiotic phenotypes and crossover rate were simply trivial consequences of the number of crossovers observed on a chromosome (**Fig. 4a**). In addition to capturing the effects of a cell or donor’s underlying meiotic proclivity rather than detected crossover number, this analysis includes the effect of any crossovers that occurred in the parent spermatocyte on the detected two-crossover chromosome’s non-observed sister chromatid. On two-crossover chromosomes, end-zone usage (**Fig. 4b**) and crossover separation (**Fig. 4c**) varied across individuals (Kruskal–Wallis chi-squared = 1,034, *df* = 19, *p* = 10^−207^ and Kruskal–Wallis chi-squared = 1,820, *df* = 19, *p* < 10^−300^, respectively) and correlated strongly and negatively with the donor’s genome-wide recombination rate (Pearson’s *r* = −0.95, *p* = 8 × 10^−11^ and Pearson’s *r* = −0.90, *p* = 5 × 10^−8^, respectively; additional control analyses described in Supplemental Text). These relationships indicate that inter-individual variation in recombination rates is a proxy for other meiotic phenotypes, including crossover interference and position preference.

**Figure 4.**
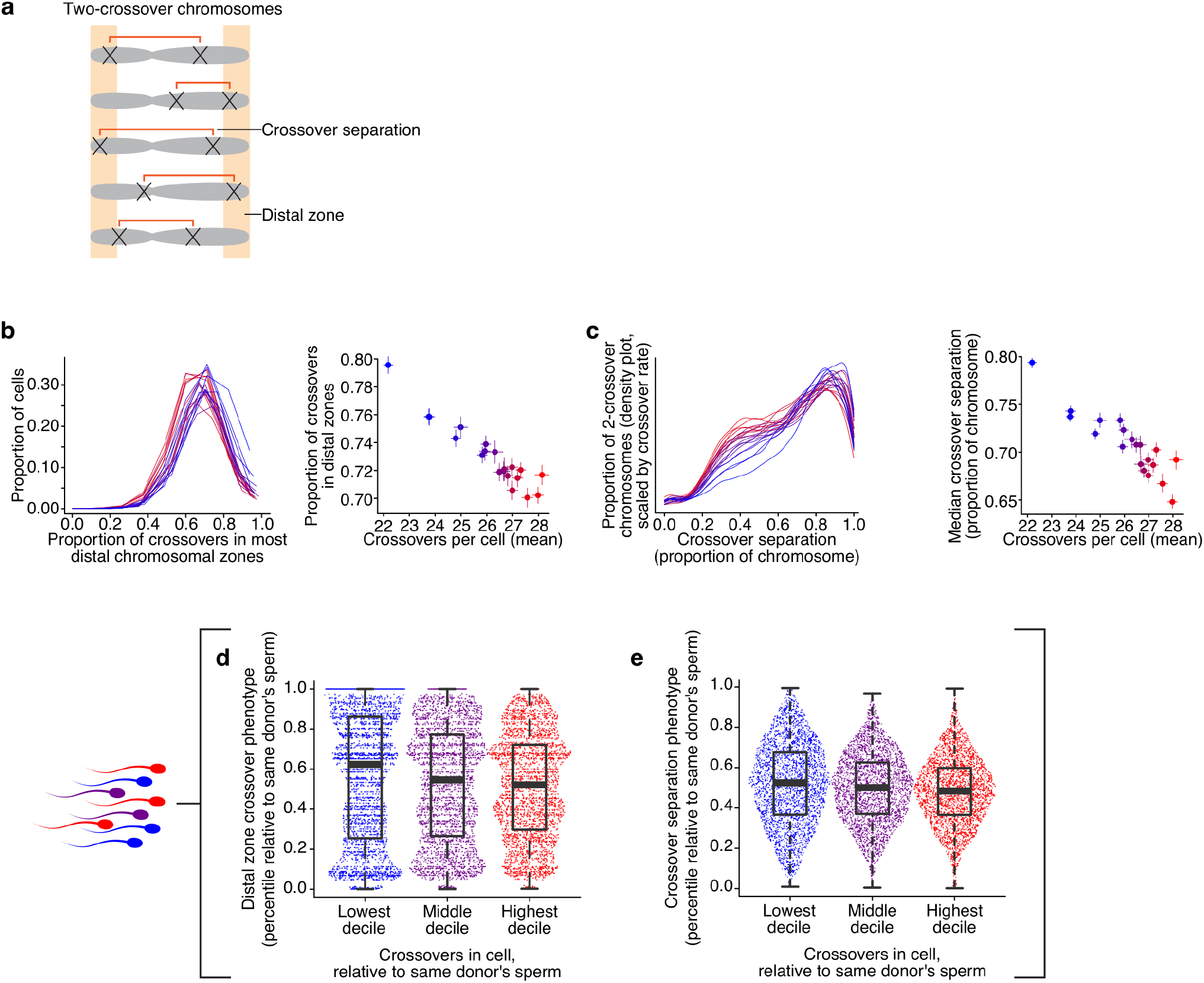
Chromosomes with two crossovers vary in crossover positioning and crossover separation (interference). **a**, Measuring the separation between crossovers, a readout of crossover interference (red brackets), and the proportion of crossovers in distal chromosome crossover zones (orange shading) on two-crossover chromosomes. The midpoint between the SNPs bounding the crossover was used as the single position for each crossover in all analyses. Error bars in (**b**) and (**c**) are 95% confidence intervals. On two-crossover chromosomes, the proportion of crossovers falling in the most distal chromosome crossover zones (**b**) and crossover separation (the distance between crossovers expressed as the proportion of the chromosome separating them) (**c**) vary among 20 sperm donors (left panels; proportion of crossovers in end per cell distributions among-donor Kruskal–Wallis chi-squared = 1,034, *df* = 19, *p* = 2 × 10^−207^; all crossover separations among-donor Kruskal–Wallis chi-squared = 1,820, *df* = 19, *p* < 10^−300^). In (**c**), density plot of separation between crossovers is shown; the area under each curve is equivalent to each donor’s global crossover rate. Right panels, both properties (y axes) shown vs. global crossover rate from all chromosomes (x axes) (correlations: proportion of all crossovers across cells’ two-crossover chromosomes in distal zones *r* = −0.95, *p* = 8 × 10^−11^; median crossover separation on two-crossover chromosomes *r* = −0.90, *p* = 5 × 10^−8^). **d**, Relationship of a cell’s distal-zone crossover phenotype (the proportion of crossovers that are in the most distal zones) to its crossover-rate phenotype; both phenotypes are analyzed as percentiles relative to other sperm from the same donor. As in (**b**), only two-crossover chromosomes were used to measure the distal zone crossover phenotype. (*n cells* per decile = 3,152, 3,080, 3,101 for first, fifth, and tenth deciles, respectively; Mann–Whitney *W* = 5,271,934.5, *p* = 2 × 10^−9^ between first and tenth deciles. Boxplots, medians and interquartile ranges; whiskers, minima to maxima. An integer effect is evident in the non-continuous distribution of the phenotype measurement because, at a single-cell level, the number of crossovers on all two-crossover chromosomes is modest.) **e**, Relationship of a cell’s crossover-separation phenotype (the median of all fractions of a chromosome separating their crossovers in each cell, measured on two-crossover chromosomes as in other panels) to its crossover-rate phenotype; both phenotypes are analyzed as percentiles relative to other sperm from the same donor. Mann–Whitney *W* = 148,548,161, *p* = 3 × 10^−53^ between first (*n* = 11,658) and tenth (*n* = 23,154) deciles (all inter-crossover separations used in test). Extended Data Fig. 11 shows more detail on crossover interference; Extended Data Figs. 12-15 show crossover separation for each autosome; Extended Data Fig. 16 shows data as in (**d**) and (**e**) for all crossover number deciles.

Single-cell analysis makes it possible to see how cellular phenotypes relate to one another, both across donors and across individual cells from the same donor. An intriguing possibility is that the same relationships generate both variation at both single-cell and person-to-person levels. To investigate this idea, we looked for connections between crossover rate and other crossover phenotypes among individual sperm cells, asking whether cells with more or fewer crossovers than the average for their donor exhibited distinct crossover interference and crossover-position-preference phenotypes. On two-crossover chromosomes, cells with more crossovers (on other chromosomes) placed a smaller fraction of their crossovers in chromosomal end zones and made their crossovers closer together (**Fig. 4d,e**, Extended Data Figure 16; Mann–Whitney *W* = 5,271,934.5; *p* = 2 × 10^−9^ in proportion of crossovers in distal zones in the 10% of cells with the highest crossover rate vs. 10% of cells with lowest crossover rate, Mann– Whitney *W* = 148,548,161, *p* = 3 ×10^−53^ result in crossover separation between cells in these same deciles of crossover rate; Methods). This result suggests that analogous relationships generate variations in meiotic outcome both among cells and across individuals (Discussion).

### Aneuploidy across chromosomes and individual sperm donors

During meiosis, a chromosome can mis-segregate (non-disjoin), yielding two aneuploid gametes in which that chromosome is reciprocally absent (a loss) or present in two copies (a gain). The frequency of paternally-derived aneuploidy is typically measured by fluorescence *in situ* hybridization (FISH) in a few chromosomes in single sperm^21–23^ or inferred genome-wide from embryos^42, 43^. We measured the ploidy of each chromosome and chromosome arm in each of the 31,228 gametes by analyzing sequence coverage (**Fig. 5a**, Methods), finding 787 whole-chromosome aneuploidies and 133 chromosome arm-scale gains and losses. All chromosomes and sperm donors were affected, with the sex chromosomes and acrocentric chromosomes (13, 14, 15, 21, and 22) having the highest rates of aneuploidy, consistent with the results of FISH studies that include chromosomes X, Y, 21, and 22^21–23, 44^ (**Fig. 5b**).

**Figure 5.**
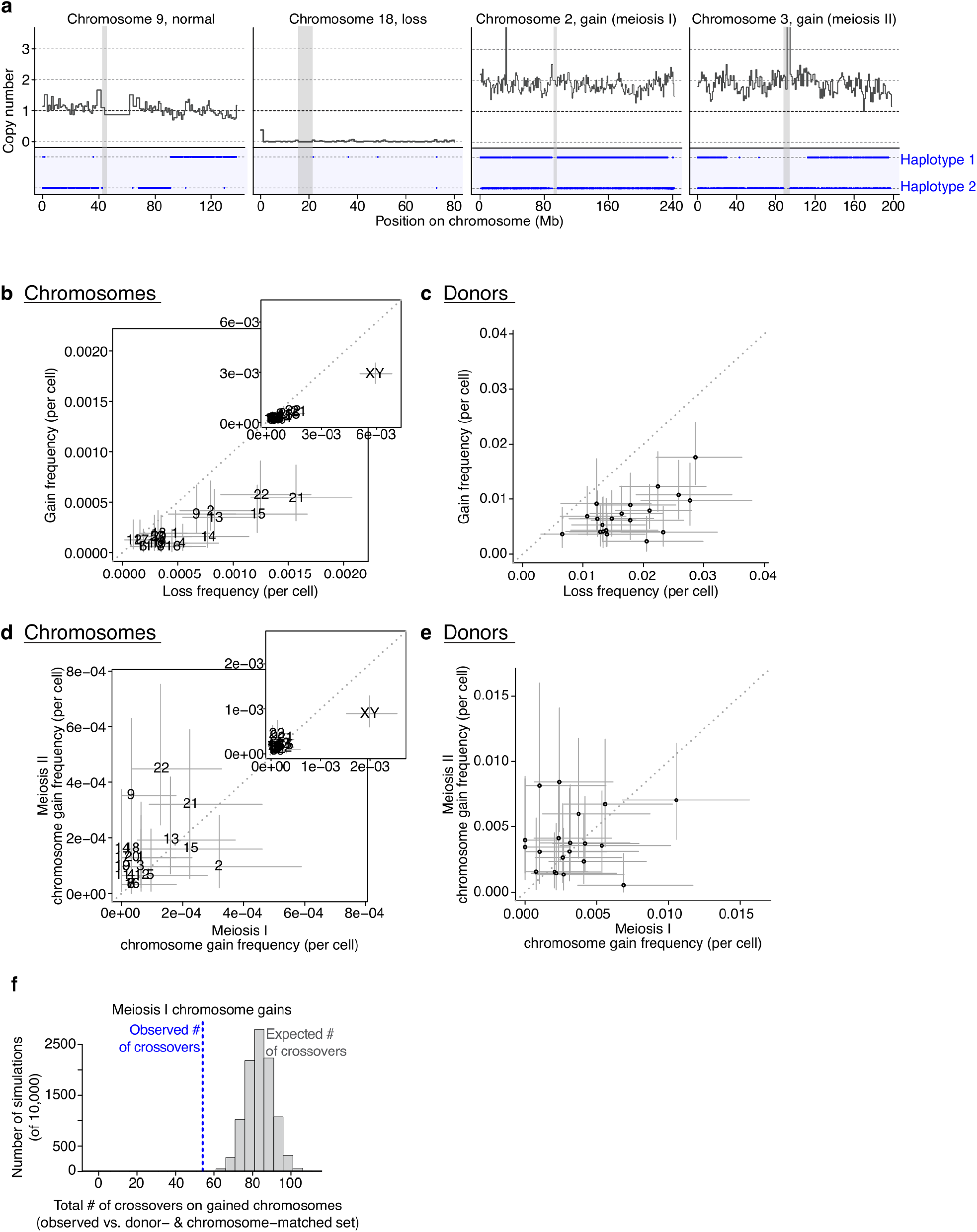
Aneuploidy in single sperm from 20 sperm donors. **a**, Example chromosomal ploidy. Copy number: thick dark gray line (normalized sequence coverage in 1 Mb bins); observed heterozygous SNP alleles: blue dots; parental haplotype of origin: dashed blue lines in bottom blue region; centromeres: gray vertical boxes. Gains occurring during nondisjunction of homologs at meiosis I (MI) have different haplotypes at their centromere (both haplotypes present, second from right). Gains occurring during nondisjunction of sister chromatids at meiosis II (MII) have identical haplotypes at their centromere (only one haplotype present, *e.g.* haplotype 2, rightmost). **b**, Frequencies of whole-chromosome losses (x axis) and gains (y axis) for each chromosome (excluding XY Pearson’s *r* = 0.88, *p* = 7 × 10^−8^; including XY [inset] Pearson’s *r* = 0.99, *p* < 10^−300^). **c**, Per-sperm-donor aneuploidy rates (axes as in **b**) (excluding XY [not shown] Pearson’s *r* = 0.51, *p* = 0.02; including XY Pearson’s *r* = 0.62, *p* = 0.003). **d**, Frequencies of whole-chromosome gains occurring during MI (x axis) and MII (y axis) for each chromosome (excluding XY Pearson’s *r* = 0.32, *p* = 0.15; including XY [inset] Pearson’s *r* = 0.85, *p* = 3 × 10^−7^). **e**, Per-sperm-donor division of origin (axes as in **d**) (excluding XY [not shown] Pearson’s *r* = 0.06, *p* = 0.80; including XY Pearson’s *r* = 0.17, *p* = 0.47). For **b-e**, error bars: binomial 95% confidence intervals on number of losses or gains divided by total number of cells (all individuals combined per chromosome, **b** and **d**; all chromosomes combined per cell, **c** and **e**; cell counts in Table 1). **f**, Total number of crossovers on MI nondisjoined chromosomes (blue line; *n* = 35 analyzed) compared to 10,000 donor- and chromosome-matched sets (35 × 2 chromosomes per set) of properly segregated chromosomes (gray histogram). (One-sided permutation *p* < 0.0001, for the hypothesis that gained chromosomes have fewer crossovers).

The frequency of aneuploidy varied 4.5-fold among individual sperm donors, who had rates of 0.010 to 0.046 aneuploidy events per cell (**Fig. 5c**, Table 1). As expected, donors with more losses also had more gains (autosomes only Pearson’s *r* = 0.51, *p* = 0.02; including XY Pearson’s *r* = 0.62, *p* = 0.003). This variation in aneuploidy rate among 20 young sperm donors (18–38 years), who were judged by clinical criteria to have healthy sperm, appears to reflect genuine inter-individual variation in vulnerability to nondisjunction (rather than statistical noise, Supplemental Text), consistent with FISH-derived observations of aneuploidy frequency in six chromosomes among 10 donors^22, 23^.

Canonically, nondisjunction creates a loss and a gain, such that one might expect sperm with chromosome losses and gains to be equally common. However, we observed 2.4-fold more losses than gains (554 losses vs. 233 gains, proportion test *p* = 2 × 10^−30^), and this asymmetry did not appear to reflect technical ascertainment bias (Supplemental Text; Extended Data Fig. 17). Among early embryos, losses of chromosomes are observed more frequently than gains, especially among paternal events^42, 43^; this imbalance has previously been attributed to post-fertilization mitotic chromosome loss, as it has not been observed in FISH studies^18, 21, 23^. However, our results suggest that gain/loss asymmetry may already be present among sperm.

Nondisjunction can occur at meiosis I (MI), when homologous chromosomes separate, or at meiosis II (MII), when sister chromatids separate. Because recombination occurs in MI (prior to disjunction) but does not occur at centromeres, homologs nondisjoined in MI will have different haplotypes at their centromeres, whereas sisters nondisjoined in MII will have the same haplotype at their centromeres (**Fig. 5a**, Methods). (On the sex chromosomes, X and Y disjoin in MI, and the sister chromatids of X and Y disjoin at MII.) Encouragingly, for chromosome 21, the principal chromosome for which earlier estimates (from patients with trisomy) were possible, our finding of 33% MI events and 67% MII events matched previous paternal estimates^45^.

Across all chromosomes, 112 gains arose during MI (50 autosomal, 62 sex chromosome) and 120 during MII (92 autosomal, 28 sex chromosome). Sex chromosomes were 2.2 times more likely to be affected in MI than MII, whereas autosomes were 2.0 times more likely to be affected in MII than MI (proportion test 35.2% MI gains on autosomes vs. 68.9% MI gains on sex chromosomes *p* = 1.3 × 10^−6^). Division-of-origin frequencies did not correlate either across chromosomes or sperm donors, implying that MI and MII have distinct nondisjunction vulnerabilities across people and individual chromosomes (**Fig. 5d,e**; across autosomes, Pearson’s *r* = 0.32, *p* = 0.15; across donors autosomes only, Pearson’s *r* = 0.06, *p* = 0.80; including XY, Pearson’s *r* = 0.17, *p* = 0.47) (consistent with studies of viable trisomies 13, 18, and 21 in embryos and individuals^45–50^).

### Relationship between aneuploidy and recombination

Although crossovers seem protective against nondisjunction in maternal meiosis^25, 48–51^, this relationship to aneuploidy is less clear in paternal meiosis^29, 45, 52–54^. To test whether nondisjunction associated with fewer crossovers in sperm, we compared the number of crossovers on gained chromosomes to those on chromosomes of normal copy number (we focused on gains because in the case of losses, it is impossible to determine what occurred on an absent chromosome). Crossovers on gained chromosomes were inferred as transitions between the presence of both haplotypes and the presence of just one haplotype. We compared the total number of crossovers on gained chromosomes (ascertainment criteria are described in Supplemental Text) to the total number of crossovers in 10,000 sets of correctly segregated chromosomes matched (to each gained chromosome) for donor and chromosome identity. Chromosome gains occurring in MI (when recombination happens) had 36% fewer total crossovers than the mean of the matched sets of well-segregated chromosomes (54 total crossovers on gains, 84.2 mean total crossovers on matched sets, one-sided permutation *p* < 0.0001), suggesting that crossovers protected against MI nondisjunction of the chromosomes on which they occurred (**Fig. 5f**). The same was not true of MII gains (Supplemental Text; Extended Data Fig. 18a).

We tested for broader relationships between crossover rates and aneuploidy at the cell and donor levels and found no clear effects, although we had limited power (Supplemental Text) (Extended Data Fig. 18b,c). One potential explanation for these findings is that the actual crossover, rather than the propensity toward crossing over in a cell or individual, is protective against aneuploidy, consistent with a model in which crossing over helps provide necessary chromosomal cohesion and/or tension for proper disjunction^55^.

### Surprising chromosome-scale genomic anomalies

Aneuploidy is thought to arise from a single nondisjunction event that leads to loss (in one gamete) or gain (in the reciprocal gamete) of one chromosome copy. Surprisingly, we detected 19 cells that had two extra copies of entire (or nearly entire) chromosomes (2, 15, 20, and 21), perhaps due to sequential nondisjunction events in MI and MII (**Fig. 6a,b**, Extended Data Fig. 19a,b). More cells had three copies of chromosome 15 (*n* = 10) than two copies of chromosome 15 (*n* = 2) (Fisher’s exact test vs. Poisson *p* = 2 × 10^−7^, Supplemental Text), raising the possibility that, for chromosome 15, MI nondisjunction leads to additional nondisjunction during MII.

**Figure 6.**
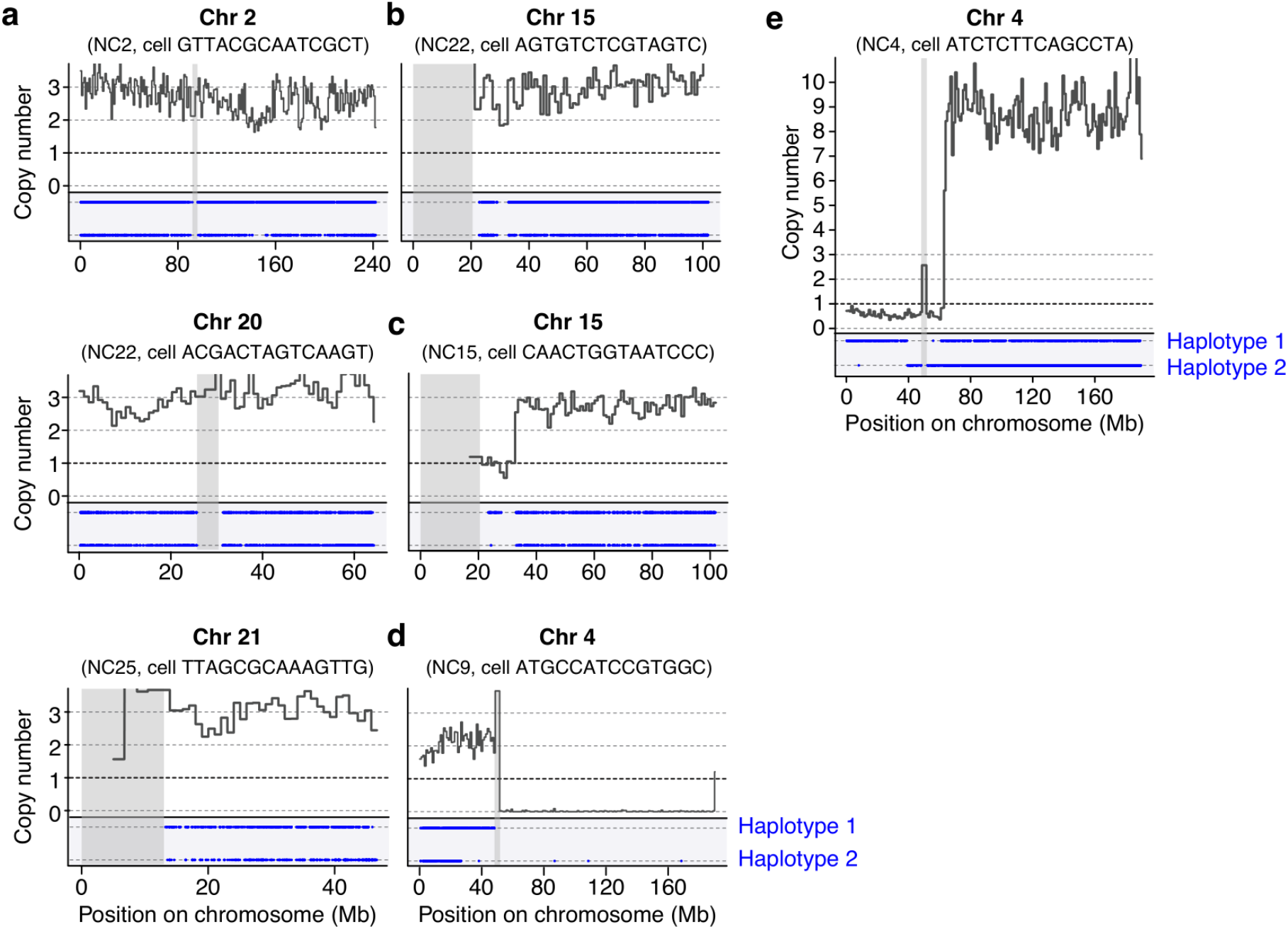
Example genomic anomalies. Copy number, SNPs, haplotypes, and centromeres (or centromeres and acrocentric arms) are as in Fig. 5a., coordinates are hg38, and donor and cell identities are noted as subtitles. Chromosomes 2, 20, 21 (**a**) and 15 (**b**) are sometimes present in an otherwise haploid sperm cell in three copies. **c**, A distinct triplication of chromosome 15, from ∼33 Mb onwards, occurs in cells from 3 donors (one shown). **d**, A compound gain of the *p* arm and loss of the *q* arm of chromosome 4. **e**, A many-copy (copy number is hard to precisely infer at high numbers) amplification of most of the *q* arm of chromosome 4 (∼127 Mb). Over-representation of this region depresses read depth in the rest of the genome to under 1. Extended Data Fig. 19 shows these and further examples.

Several sperm had complex aneuploidy events that were not explained by nondisjunction. These included: multiple cells with three copies of most, but not all, of the *q* arm of chromosome 15; one cell that gained the *p* arm of chromosome 4 while losing the *q* arm; and one cell with at least eight copies of most of the *q* arm of chromosome 4 (**Fig. 6c-e;** Extended Data Fig. 19c,d). We estimate that the gamete with at least eight copies of 127 Mb of 4*q* contained a minimum of 890 Mb of extra genomic DNA, demonstrating that the human sperm nucleus can accommodate at least 30% more DNA than is typically in the haploid genome (**Fig. 6e**). This gamete carried both parental haplotypes of chromosome 4, though the extra copies came from just one of the two parental haplotypes (93% of observed alleles of heterozygous SNPs in the amplified region were haplotype 2). We know of no mechanism that would generate such a gamete.

## Discussion

The genomes of 31,228 human sperm cells revealed interconnected variation among diverse meiotic phenotypes. These relationships existed at different and sometimes multiple levels: (i) individuals’ average meiotic phenotypes; (ii) variation among single sperm cells from the same person; and (iii) specific chromosomes and events.

Rates of aneuploidy varied conspicuously (from 1.0% to 4.6%) among the 20 young sperm donors (**Fig. 5c**). Aneuploidy was less likely when a chromosome had more crossovers, though at higher levels of organization (cells and donors) aneuploidy rates and crossover rates varied independently (Extended Data Fig. 18). Some chromosomes were more vulnerable to nondisjunction in MI, and others to nondisjunction in MII; some donors were more vulnerable to nondisjunction in MI, and others to nondisjunction in MII (**Fig. 5d,e**). These results suggest a complex landscape of vulnerability to aneuploidy in which inter-individual variation is multi-faceted and considerable in magnitude.

Inter-individual variation in crossover rates has previously been visible through computational analyses of SNP data^1–10^. Here, single-gamete sequencing revealed that donors with high crossover rates also exhibit other meiotic phenotypes, including a tendency to make crossovers closer together and to place a smaller fraction of their crossovers in telomere-proximal zones (**Figs. 3**, **4**). The same underlying biological variation may shape all three phenotypes (rate, location, and separation).

Individual cells from the same donor also appeared to have underlying meiotic proclivities that coordinated these meiotic outcomes across the genome and with one another. This was observed in the correlation of crossover number across different chromosomes: even among cells from the same donor, gametes with more crossovers in half of their genome tended to have more crossovers in the other half. High-crossover-rate cells also made pairs of crossovers (on the same chromosome) closer together (in genomic distance) and placed proportionally fewer of their crossovers in telomere-proximal chromosomal regions (**Fig. 4d,e**).

What could cause these meiotic phenotypes to be coupled to one another, across chromosomes and at multiple levels of organization (cells and individuals)? Intriguingly in this regard, the physical length of meiotic chromosomes – which is inversely related to their degree of compaction – has been observed to vary among meiotic cells, and individual cells with more compacted (shorter) chromosomes also tend to have fewer crossovers^24–26, 56^. A simple model (**Fig. 7**) might explain the observed correlations: cell-to-cell and person-to-person variation in the compaction of meiotic chromosomes could cause the variation in and correlations among crossover rate, location, and interference, provided that crossover interference occurs as a function of physical (micron) distance along the meiotic chromosome axis/synaptonemal complex rather than genomic (base pair) distance^25, 34, 57, 58^ and that the first crossover on a chromosome is more likely to occur near a telomere^11–13^ (**Fig. 7**).

**Figure 7.**
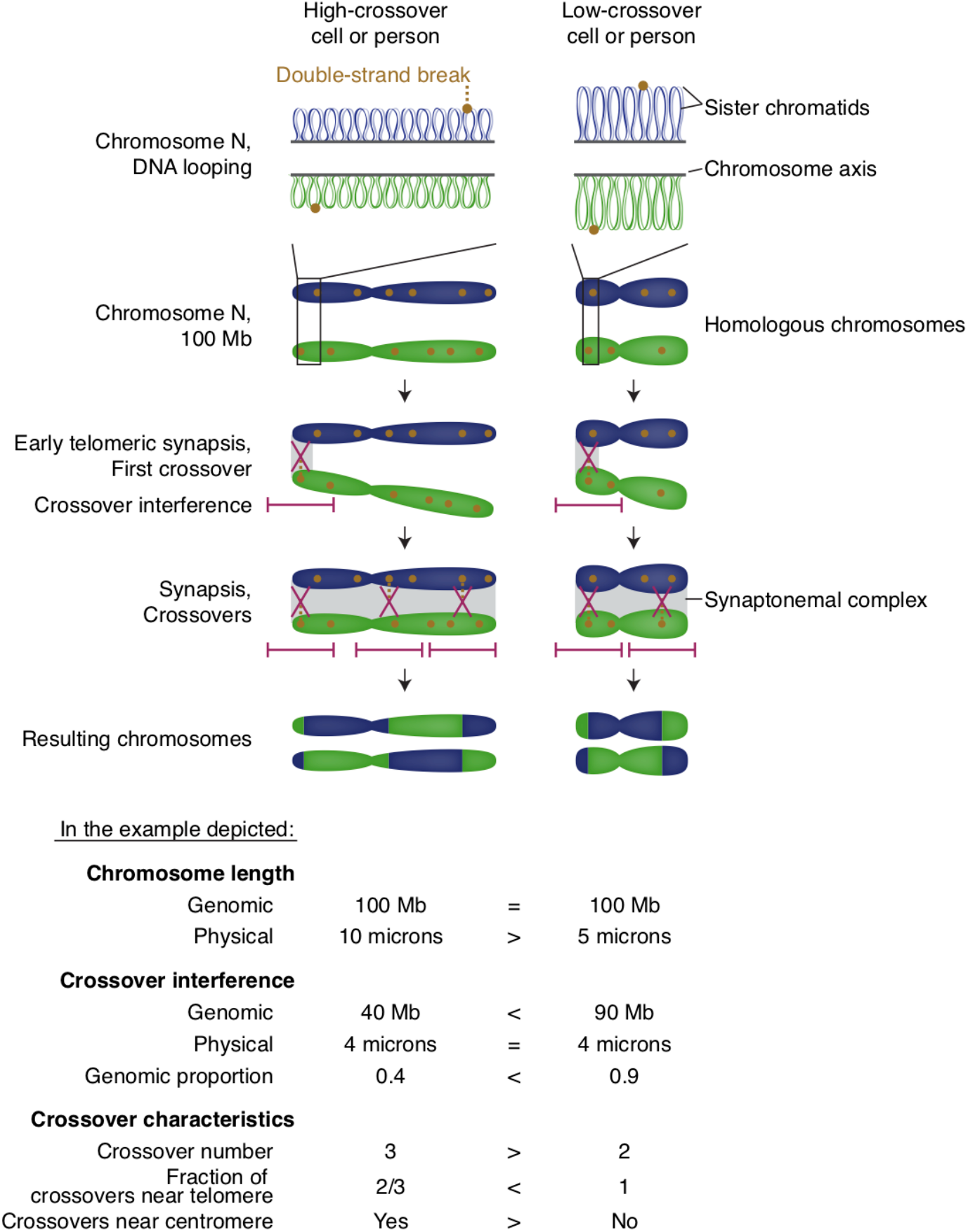
Meiotic phenotype variation among single gametes and individuals may be governed by variation in the physical compaction of chromosomes. The physical length of the same chromosome varies among spermatocytes at the pachytene stage of meiosis, likely by differential looping of DNA along the meiotic chromosome axis (*e.g.* left column shows smaller loops, resulting in more loops total and in greater total axis length compared to the right column with larger loops)^13,59–62^. This physical chromosome length is correlated across chromosomes among cells from the same individual^25, 26^ and correlates with crossover number^13, 24–26, 56, 60^. This length – measured as the length of the chromosome axis or of the synaptonemal complex (the connector of homologous chromosomes) – can vary two or more-fold among a human’s spermatocytes^26^. We propose that the same process differs on average across individuals and partially explains inter-individual variation in recombination rate: on average, individual 1 (left) would have meiotic chromosomes that are physically longer (less compacted) in an average cell than individual 2 (right); one example chromosome is shown in the figure. After the first crossover on a chromosome (likely at the telomere, where synapsis typically begins in male human meiosis before spreading across the whole chromosome^11–13^), crossover interference prevents nearby double-strand breaks from becoming crossovers; double-strand breaks far away can become crossovers (which themselves also cause interference). Crossover interference occurs over relatively fixed physical (micron) distances^25, 34, 57, 58^, but these distances encompass different genomic (Mb) amounts of DNA and therefore proportions of the chromosome when meiotic chromosomes are of different lengths due to variable compaction. Thus, interference tends to lead to different total number of crossovers as a function of degree of compaction, with the resulting negative relationship of crossover interference (measured in base pairs) with crossover rate. Given that the first crossover likely occurs near the telomere, this model can also explain the negative correlation of rate and the proportion of crossovers in chromosome ends: a second crossover can only occur in a centromeric region on a chromosome that is physically long enough for interference not to block crossovers closer to the centromere. *Note: this figure shows the total number of crossovers, crossover interference extent, and crossover locations for both sister chromatids of each homolog combined; in reality, these crossovers are distributed among both sister chromatids (such that these relationships are harder to detect in daughter sperm cells, requiring large numbers of observations)*.

Human genetics research has revealed that recombination phenotypes are heritable and associate with common SNPs at many genomic loci^4, 6–10^. The largest genome-wide association study of crossover phenotypes recently found that variation in crossover rate and placement is associated with SNP haplotypes near genes that encode components of the synaptonemal complex, which connects and compacts meiotic chromosomes^8^. It is reasonable to hypothesize that inherited genetic variation at these loci might bias the average degree of compaction along the chromosome axis or synaptonemal complex, particularly given that this same property varies among cells from the same donor^24–26^. Such a model would offer a natural integration of observations about inter-individual and gamete-to-gamete variation, and of relationships among diverse meiotic phenotypes (**Fig. 7**).

Our results suggest that, in meiosis, a shared set of patterns and constraints shapes inter- and intra-individual (single-cell) variation in meiotic outcomes. It is an intriguing possibility that such parallel relationships manifest in diverse aspects of cellular biology and genetics.

## Acknowledgments

We thank Giulio Genovese for suggestions on analyses, Evan Macosko for advice on technology development, and other members of the McCarroll lab, including Chris Whelan, Steven Burger, and Bob Handsaker for their advice. We thank Mark Daly, Joel Hirschhorn, Stephen Elledge, and Samantha Schilit for their insights, 10X Genomics for discussions about reagents, and Christina L. Usher and Christopher K. Patil for contributions to the manuscript text and figures. This work was supported by R01 HG006855 to S.A.M., by a Broad Institute NextGen award to S.A.M., and by a Harvard Medical School Program in Genetics and Genomics NIH Ruth L. Kirchstein training grant to A.D.B.

## Author Contributions

A.D.B. and S.A.M. conceived and led the studies. A.D.B, S.A.M., and C.J.M. developed the experimental methods. A.D.B. and C.J.M. performed the experiments and generated the data. A.D.B and S.A.M. designed the analysis strategies, and A.D.B. performed the analyses. A.D.B., J.N., S.A.B., and A.W. wrote the software and analytical methods. A.D.B. and S.A.M. wrote the manuscript with contributions from all authors.

## Data Availability

Crossover and aneuploidy data (individual events and counts per donor and/or cell) are available via Zenodo, http://dx.doi.org/10.5281/zenodo.2581571. Raw sequence data will be deposited in the SRA via dbGaP (in process).

## Code Availability

Analysis scripts and documentation are available via Zenodo, http://dx.doi.org/10.5281/zenodo.2581596.

**Supplementary Information**, which includes Supplemental Text and Methods, is available.

**Extended Data Figures** follow in this document.

## Extended Data Figures

**Extended Data Figure 1.**
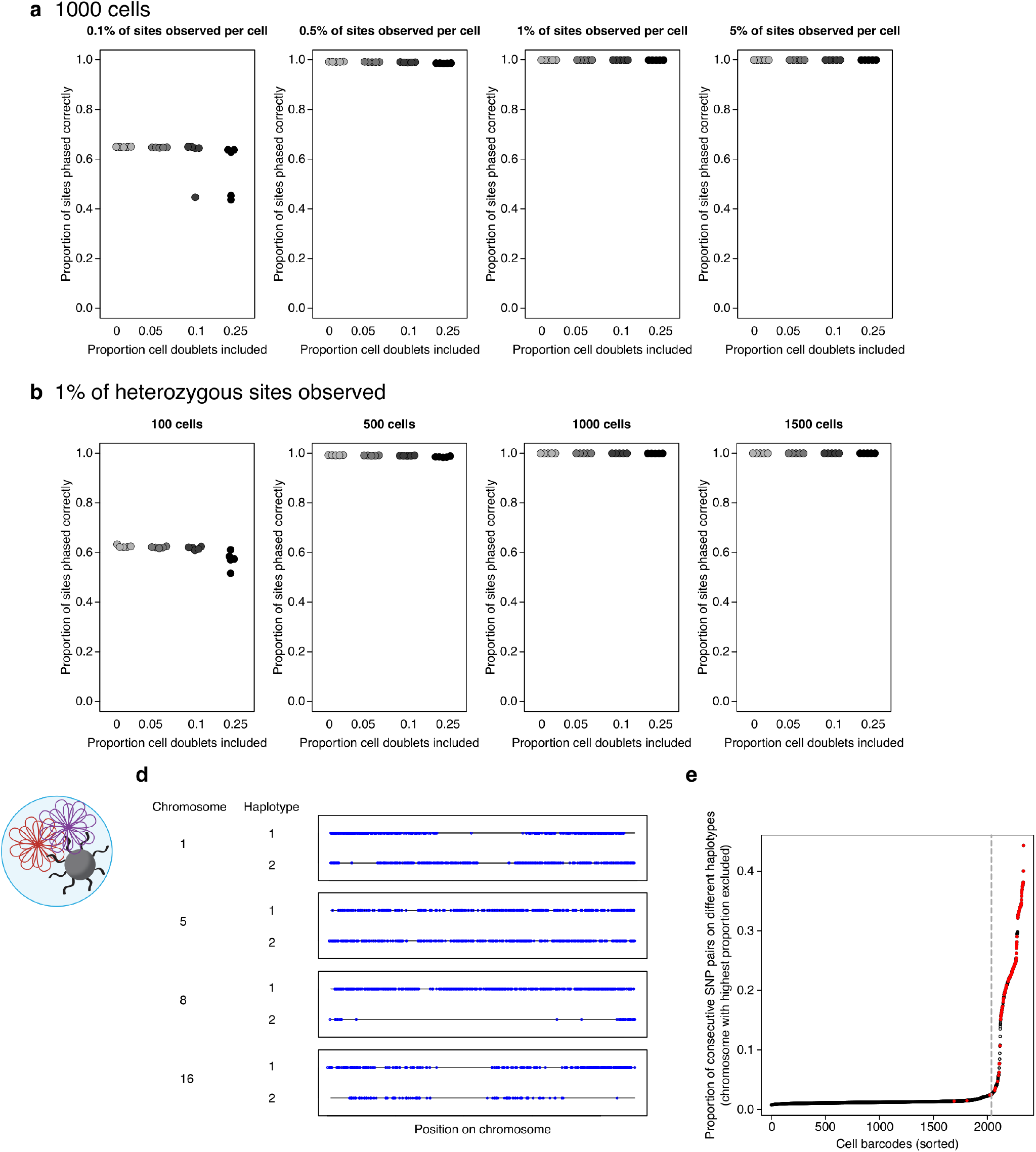
Evaluation of chromosomal phasing and identification of cell doublets. **a**, Evaluation of our phasing method (using HapCUT^63, 64^) using 1,000 simulated single-sperm genomes (generated from two *a priori* known parental haplotypes and sampled at various levels of coverage as shown in the three plots). Since cell doublets (which combine two haploid genomes and potentially two haplotypes at any region) can in principle undermine phasing inference, we included cell doublets in the simulation (in proportions shown on the X axis, which bracket the observed doublet rates). Each point shows the proportion of SNPs phased concordantly with the correct (*a priori* known) haplotypes (Y axis) for one simulation (five simulations were performed per proportion of cell doublets-percentage of observed sites condition pair). **b**, Relationship of phasing capability to number of cells analyzed. Data are as in (**a**), but for different numbers of simulated cells, all simulations with an among-cell mean of 1% of heterozygous sites observed. **c**, A cell doublet: when two cells are co-encapsulated in the same droplet, their genomic sequences will be tagged with the same barcode; such events must be recognized computationally and excluded from downstream analyses. **d**, Four example chromosomes from a cell barcode associated with two sperm cells (a cell doublet). Black lines: haplotypes; blue circles: observations of alleles, shown on the haplotype from which they derive. Both parental haplotypes are present across regions of chromosomes where the cells inherited different haplotypes. **e**, Computational recognition of cell doublets in Sperm-seq data (from an individual sperm donor, NC11). The proportion of consecutively observed SNP alleles derived from different parental haplotypes is used to identify cell doublets; this proportion is generally small (arising from sparse crossovers, PCR/sequencing errors, and/or ambient DNA) but is much higher when the analyzed sequence comes from a mixture of two distinct haploid genomes (of which, on average, half will derive from distinct parental haplotypes). We use 21 of the 22 autosomes to calculate this proportion, excluding the autosome with the highest such proportion given the possibility that a chromosome is aneuploid. The dashed gray line marks the inflection point beyond which sperm genomes are flagged as potential doublets and excluded from downstream analysis. Red points indicate barcodes with coverage of both the X and Y chromosome (potentially X+Y cell doublets or XY aneuploid cells); black points indicate barcodes with one sex chromosome detected (X or Y). The red (XY) cells below the doublet threshold are XY aneuploid but appear to have just one copy of each autosome.

**Extended Data Figure 2.**
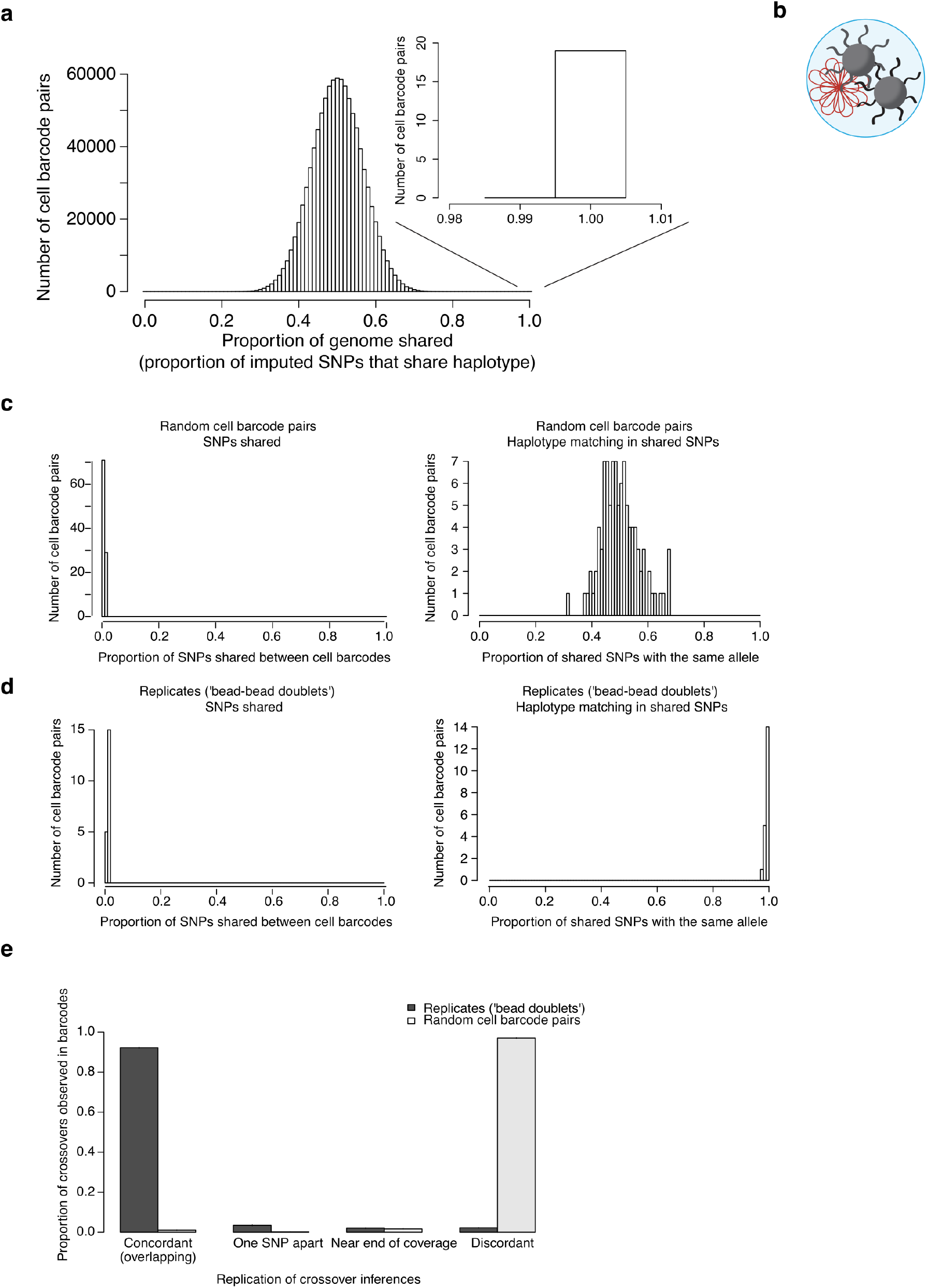
Identification and use of “bead doublets.” **a**, SNP alleles were inferred genome-wide (for each sperm genome) from its Sperm-seq data and the Sperm-seq-inferred parental haplotypes; for each pair of sperm genomes (cell barcodes), the proportion of all SNPs at which they shared the same imputed allele was estimated. A small but extremely surprising number of such pairwise comparisons (19 of 984,906 from the donor shown, NC14) indicate essentially identical genomes. **b**, We hypothesize that this arises from a heretofore undescribed scenario we call “bead doublets”, in which two barcoded beads have been co-encapsulated with the same gamete and whose barcodes therefore tagged the same haploid genome. **c**, Random pairs of cell barcodes (here 100 pairs selected from donor NC10) tend to interrogate few of the same SNPs (left), and to detect the same parental haplotype on average at the expected 50% of the genome (right). **d, “**Bead doublet” barcode pairs (here 20 pairs from donor NC10, who had the median number of bead doublets) also interrogate few of the same SNPs, yet detect identical haplotypes throughout the genome. Results were consistent across donors. **e**, Use of “bead doublets” to characterize the concordance of crossover inferences between distinct samplings of the same haploid genome by different barcodes. Analyses of the bead doublets (barcode pairs) were compared to 100 random barcode pairs per donor. Crossover inferences were classified as “concordant” (overlapping, detected in both barcodes), as “one SNP apart” (separated by just one SNP, detected in both barcodes), as “near end of coverage” (within 15 heterozygous SNPs of the end of SNP coverage at a telomere, where power to infer crossovers is partial), or as discordant. Error bars (with small magnitude) show binomial 95% confidence intervals for the number of crossovers per category divided by number of crossovers total in both barcodes (32,714 crossovers total in 1,201 bead doublet pairs; 67,862 crossovers total in 2,000 random barcode pairs; some barcodes are in multiple bead doublet or random barcode pairs).

**Extended Data Figure 3.**
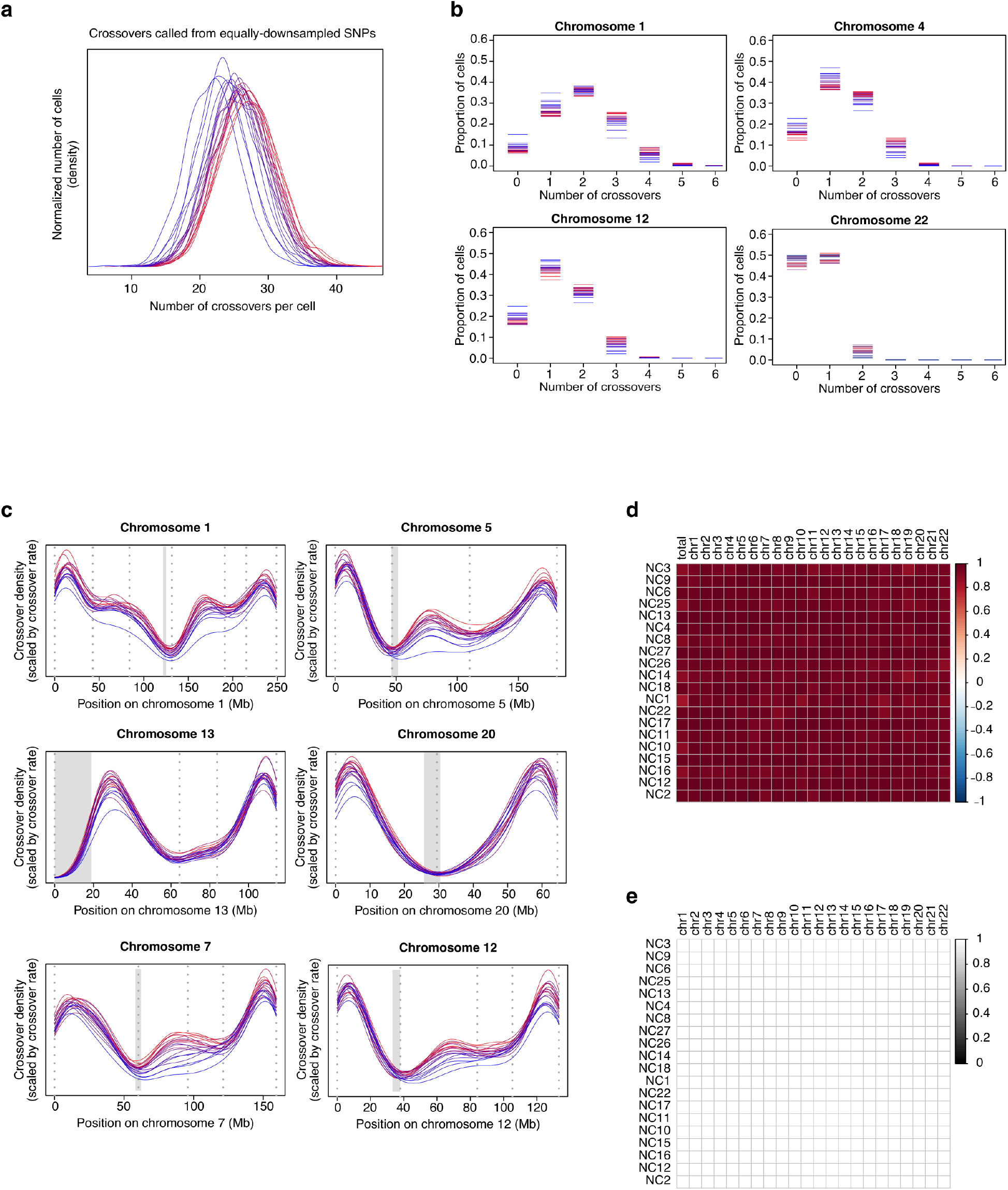
Numbers and locations of crossovers called from down-sampled data (randomly chosen equal number of SNPs in each cell). To eliminate any potential effect of unequal sequence coverage across donors and cells, down-sampling was used to create data sets with equal numbers of heterozygous SNPs typed in each cell. Crossovers were called from these random equally sized sets of SNPs from all cells. **a** and **b**, Crossover number per cell globally (**a**) and per chromosome (**b**) (as in **Fig. 2b,c**; 785,476 total autosomal crossovers called from down-sampled SNPs included, 30,778 cells included, aneuploid chromosomes excluded). **c**, (As in **Fig 3b**.) Density plots of crossover location with crossover midpoints plotted and area scaled to be equal to per-chromosome crossover rate. Gray rectangles mark centromeric regions; hg38. **d**, Similar numbers of crossovers were called from full data and equally down-sampled SNP data: we performed correlation tests for each donor and chromosome to compare the number of crossovers called from all data to the number of crossovers called from equal numbers of randomly down-sampled SNPs. Each row is a donor and each column is a chromosome (except the first column, which is global crossover number); color corresponds to Pearson’s *r* value (all chromosome comparisons *r* > 0.86, all *p* < 10^−300^). **e**, Crossovers called from equally down-sampled SNP data were in similar locations to those called from all data: we used Kruskal–Wallis tests to compare the distribution of crossover location of crossovers called from all data to the distribution of crossover location of crossovers called from equally down-sampled SNP data. Each row is a donor and each column is a chromosome; color corresponds to the Kruskal–Wallis *p* value (all *df* = 1, *p* = 1).

**Extended Data Figure 4.**
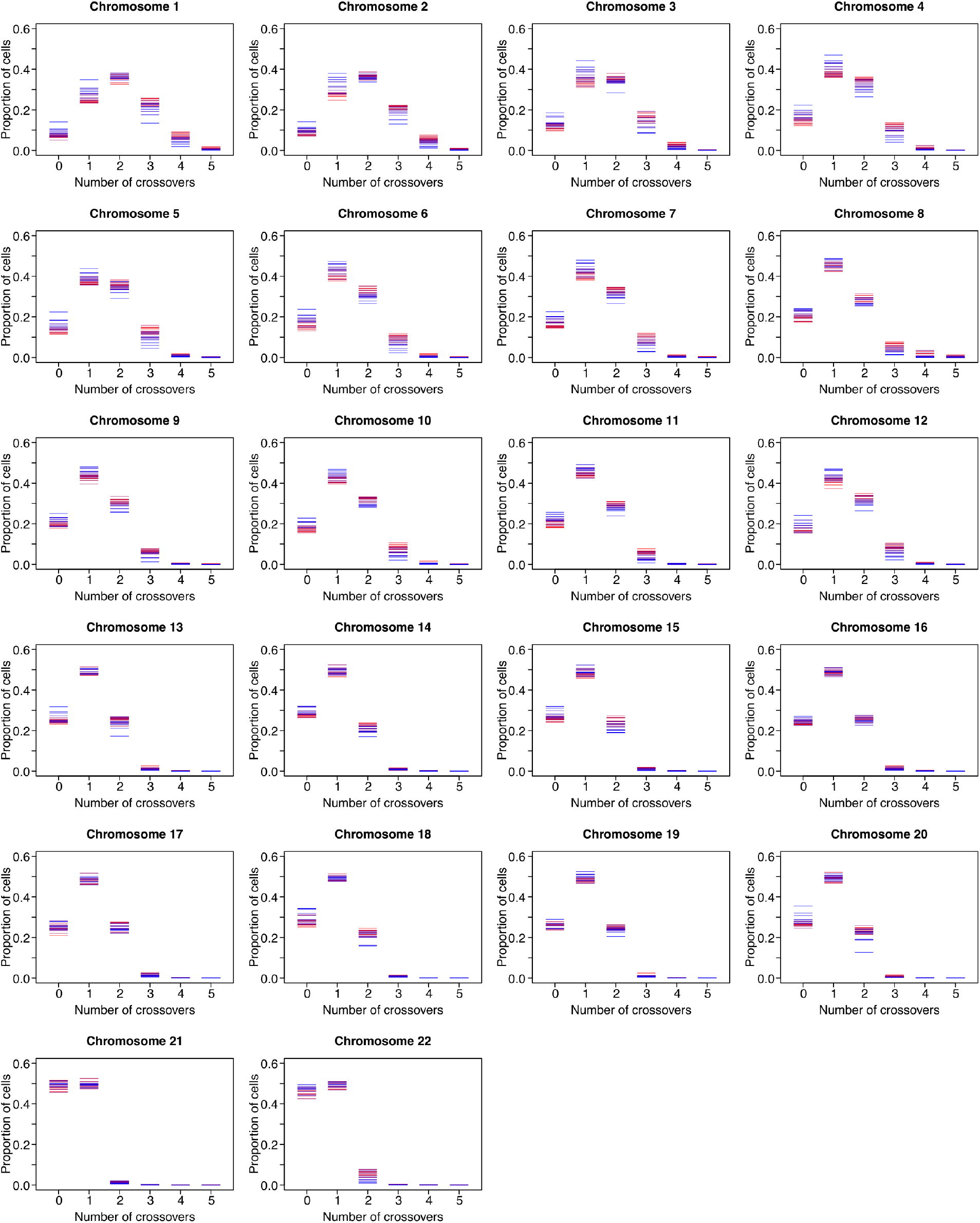
**Per-chromosome crossover** count for 20 sperm donors (colored blue [low] to red [high] based on global average crossover rate) and 22 autosomes showing the proportion of sperm cells with each crossover number per chromosome (aneuploid chromosomes excluded from crossover calling), as in **Fig. 2c**.

**Extended Data Figure 5.**
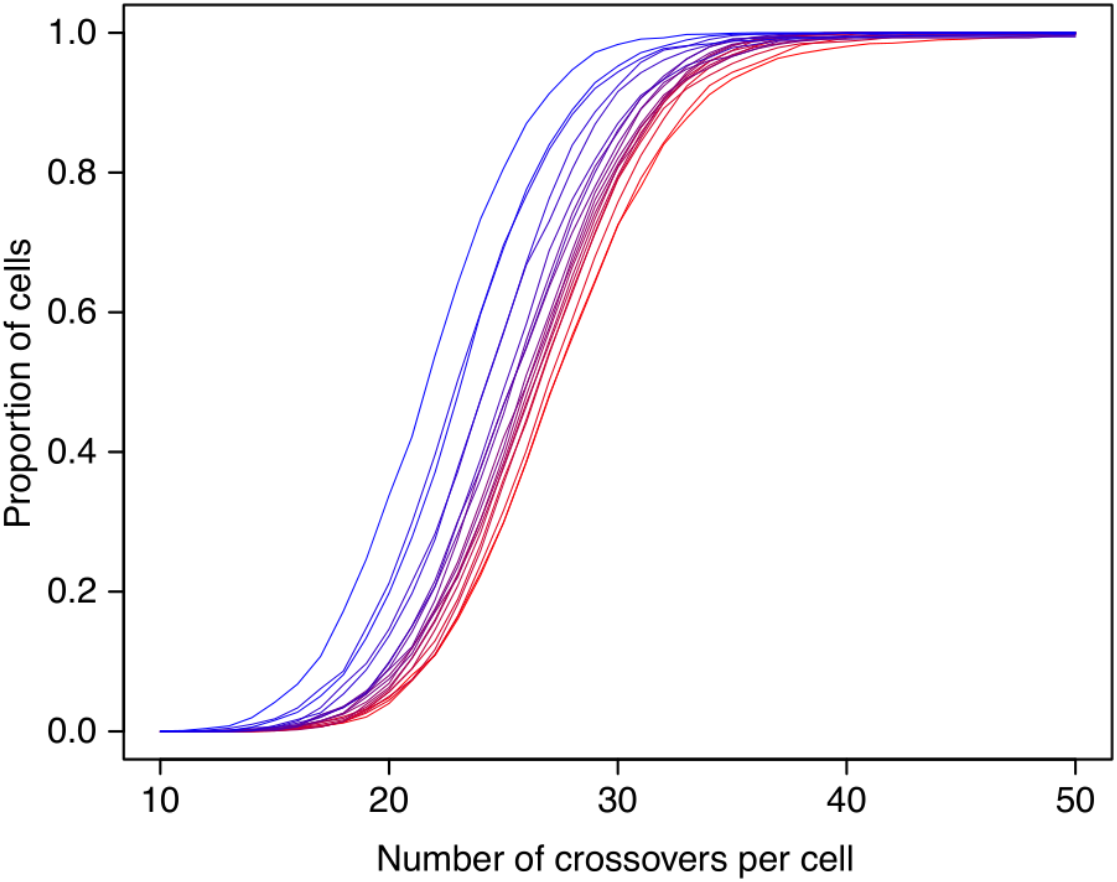
**Cumulative distribution of number of crossovers per cell** across 20 sperm donors (color corresponds to mean crossover rate, all 813,122 autosomal crossovers shown [aneuploid chromosomes excluded from crossover calling]). All 31,228 cells are included (same data as in **Fig. 2b**).

**Extended Data Figure 6.**
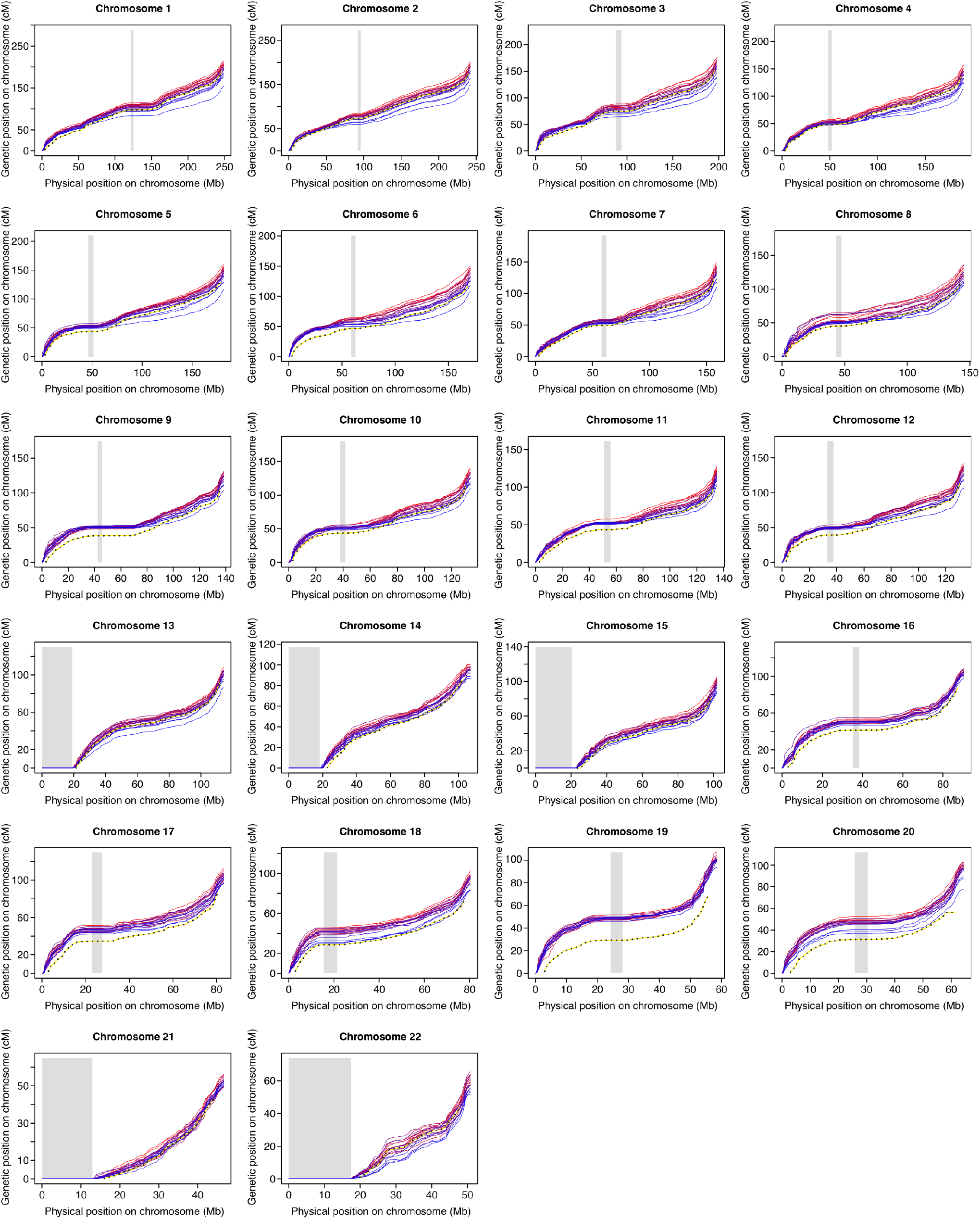
**Per-chromosome individualized genetic maps** for 20 sperm donors and 22 autosomes, as in **Fig. 2e** (dashed line highlighted with yellow, deCODE’s^10^ paternal pedigree-based genetic map, which excludes SNPs within 2.5 Mb of the most telomeric SNP observed). Physical vs. genetic distances plotted at 500 kb intervals (hg38). Gray boxes denote centromeric regions (or centromeres and acrocentric arms).

**Extended Data Figure 7.**
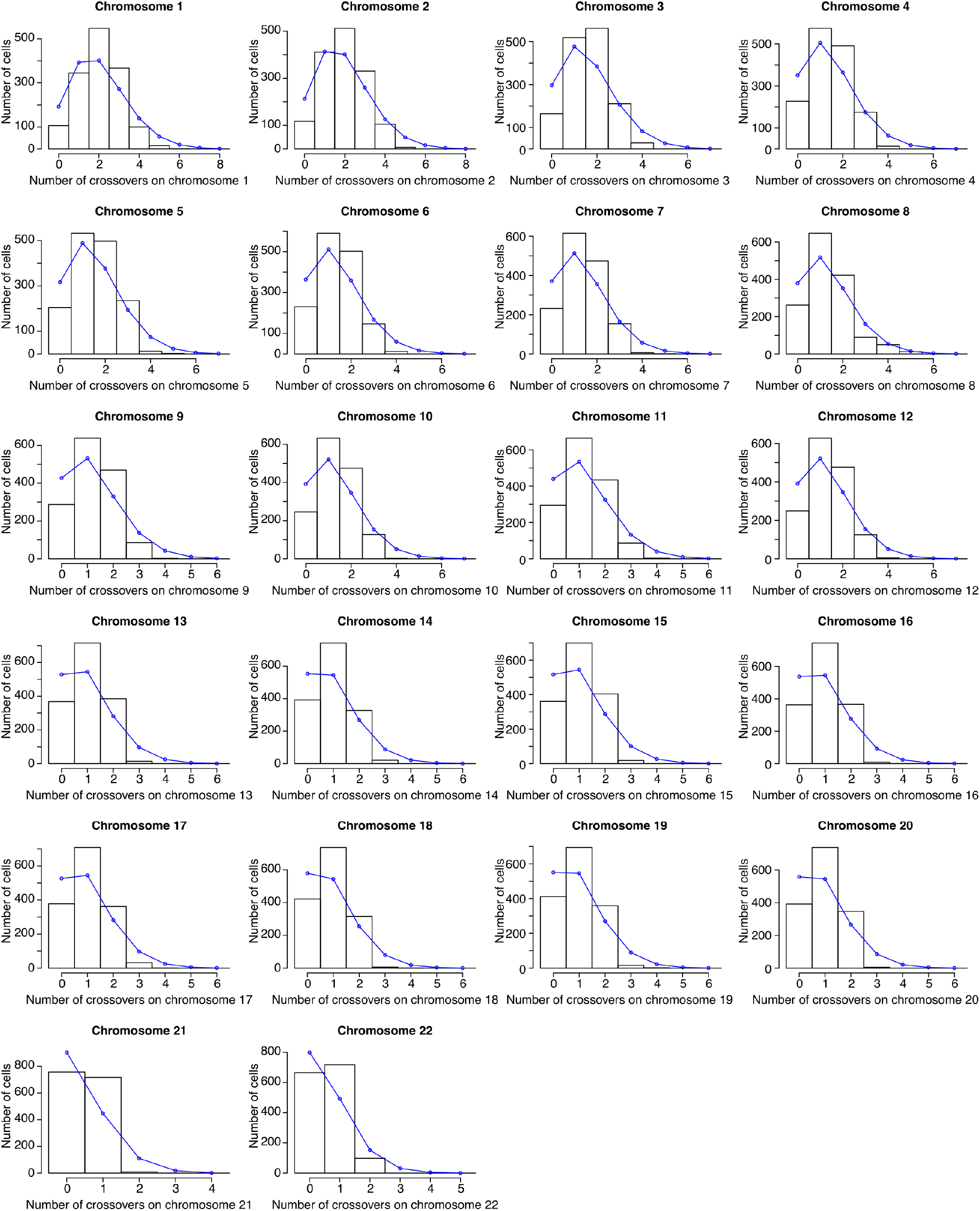
Observed versus randomly expected number of crossovers per chromosome. Random (Poisson) expectation of the number of cells with each number of crossovers on each chromosome, blue line; observed crossover number distribution, black histogram. Data shown is from donor NC4, median donor in this analysis (overall 10^th^ of 20 in both significance vs. Poisson in a chi-squared test of goodness of fit and 10^th^ of 20 in expected Poisson vs. observed chi-squared test of variance for total crossover number). All chromosomes’ distributions are significantly different from Poisson (least significant chromosome in any donor *p* = 1 × 10^−9^).

**Extended Data Figure 8.**
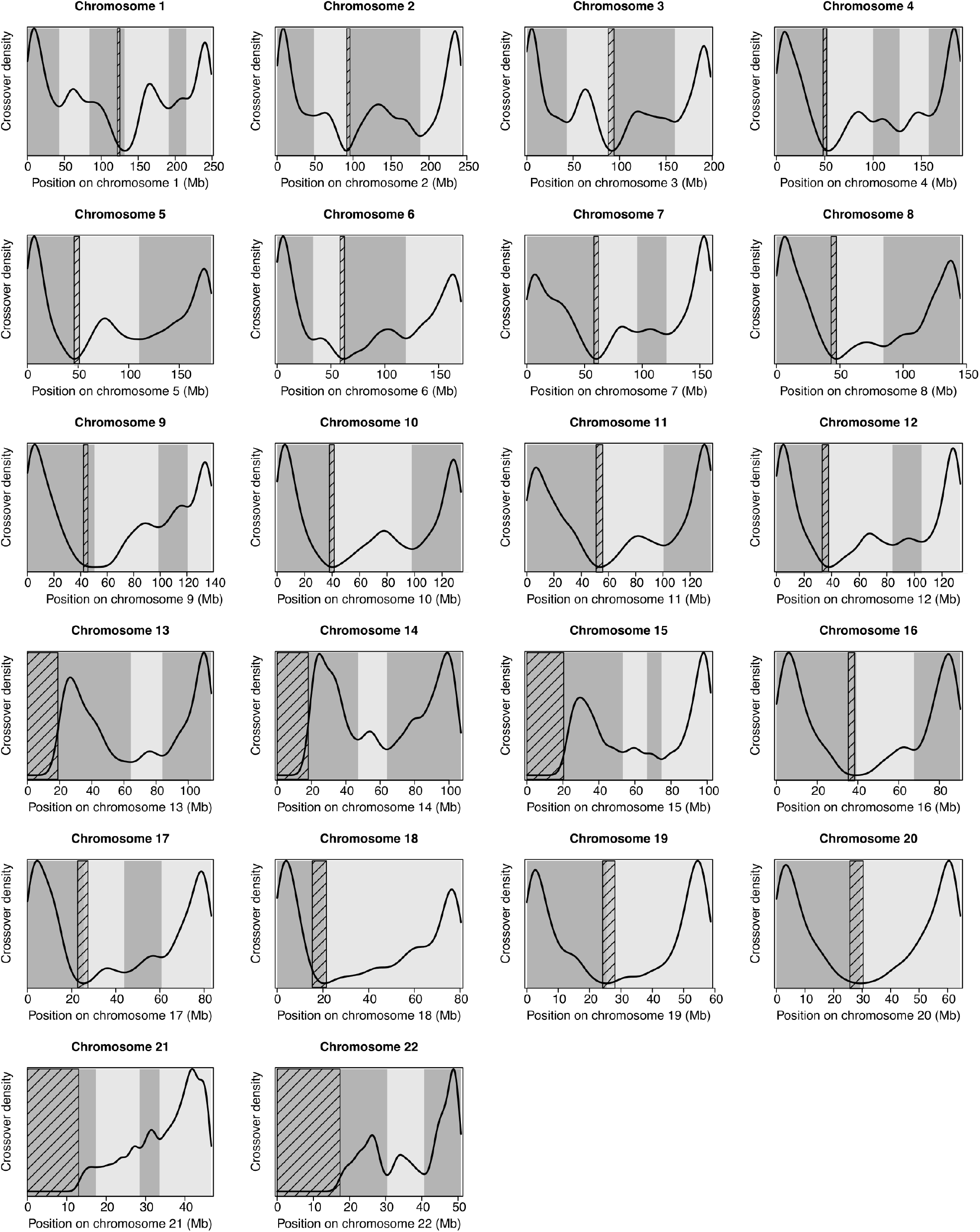
**Identification of chromosomal zones of recombination use (“crossover zones”)** from all donors’ crossovers for 22 autosomes. Density plots of crossover location for all sperm donors’ total 813,122 crossovers (aneuploid chromosomes excluded; crossover location is the midpoint between SNPs bounding crossovers) along autosomes (hg38) are shown. Crossover zones, alternating shaded and unshaded chromosomal regions. Local minima of crossover density functions mark their boundaries. Diagonally-hatched rectangles, centromeres (or centromeres and acrocentric arms).

**Extended Data Figure 9.**
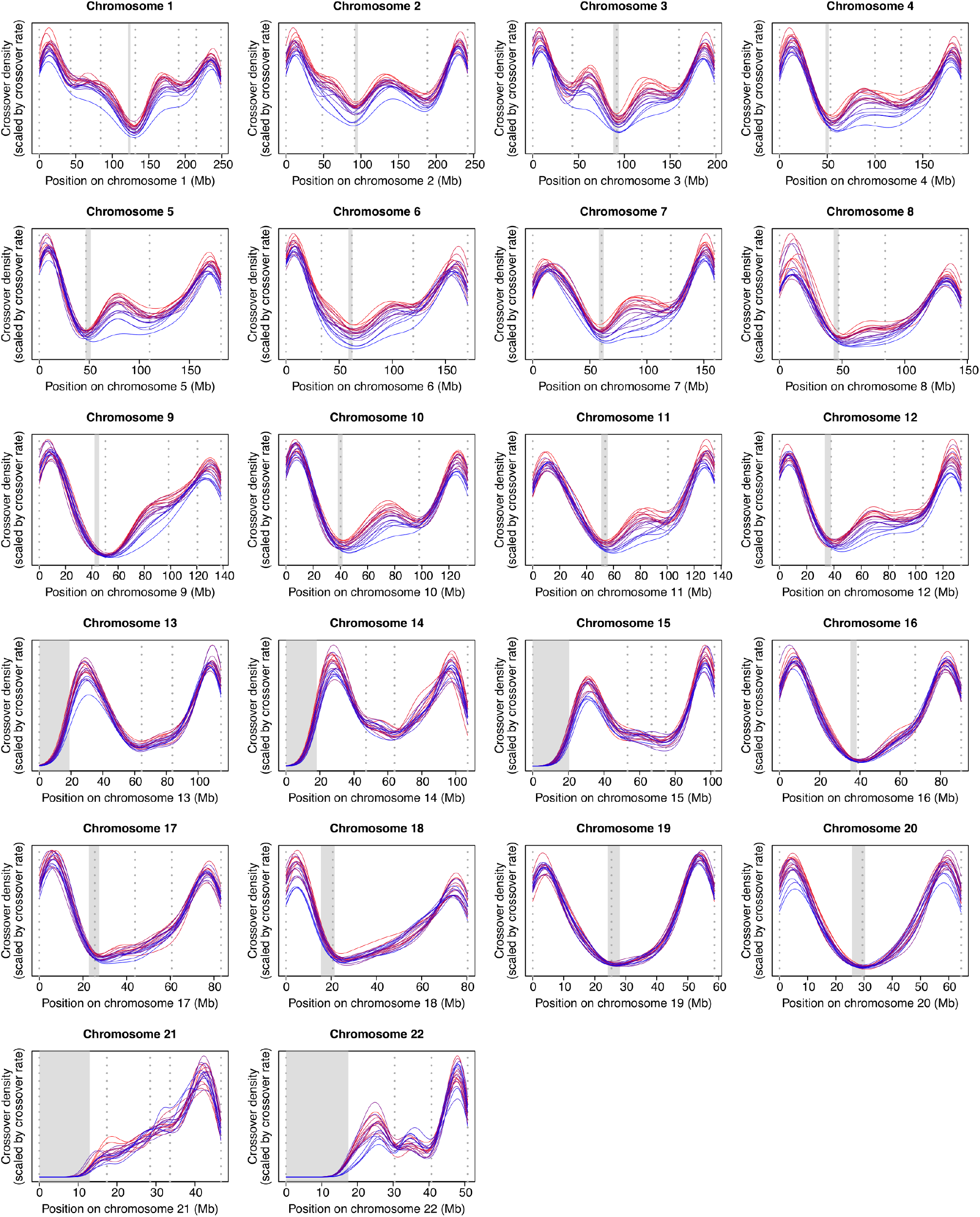
**Crossover location density plots** (normalized to crossover rate) for 20 sperm donors for 22 autosomes, as in **Fig. 3b**. The area under each curve is equivalent to the crossover rate on that chromosome for each donor. Dotted gray vertical lines denote crossover zone boundaries; gray boxes mark centromeres (or centromeres and acrocentric arms). Coordinates are in hg38.

**Extended Data Figure 10.**
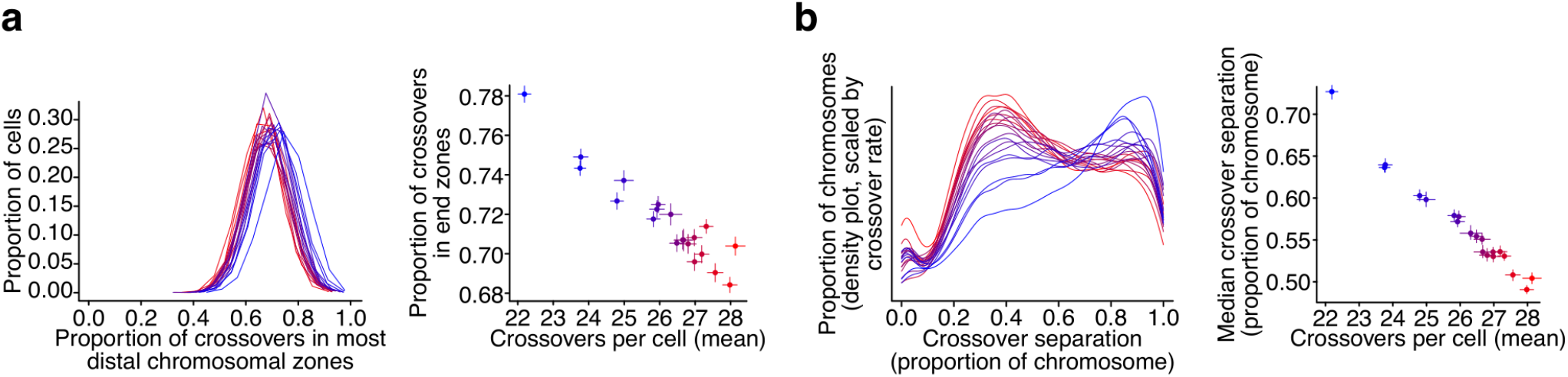
Crossover placement in end zones and crossover separation vary and correlate with crossover rate among sperm donors (all cells, chromosomes included). The midpoint between the SNPs bounding the crossover was used as the single position for each crossover in all analyses. The proportion of crossovers falling in the most distal chromosome crossover zones (a) and crossover separation, a readout of crossover interference, the distance between consecutive crossovers expressed as the proportion of the chromosome separating them (b) vary among 20 sperm donors (left panels; proportion of crossovers in end per cell distributions among-donor Kruskal–Wallis chi-squared = 2,334, *df* = 19, *p* < 10^−300^; all distances between consecutive crossovers among-donor Kruskal–Wallis chi-squared = 4,316, *df* = 19, *p* < 10^−300^). Right panels show both properties (y axes) vs. donor’s global crossover rate (x axes) (Correlation results for 20 sperm donors: proportion of all crossovers across cells in end *r* = −0.95, *p* =2 × 10^−10^; median distance between consecutive crossovers *r* = −0.99, *p* = 9 × 10^−16^).

**Extended Data Figure 11.**
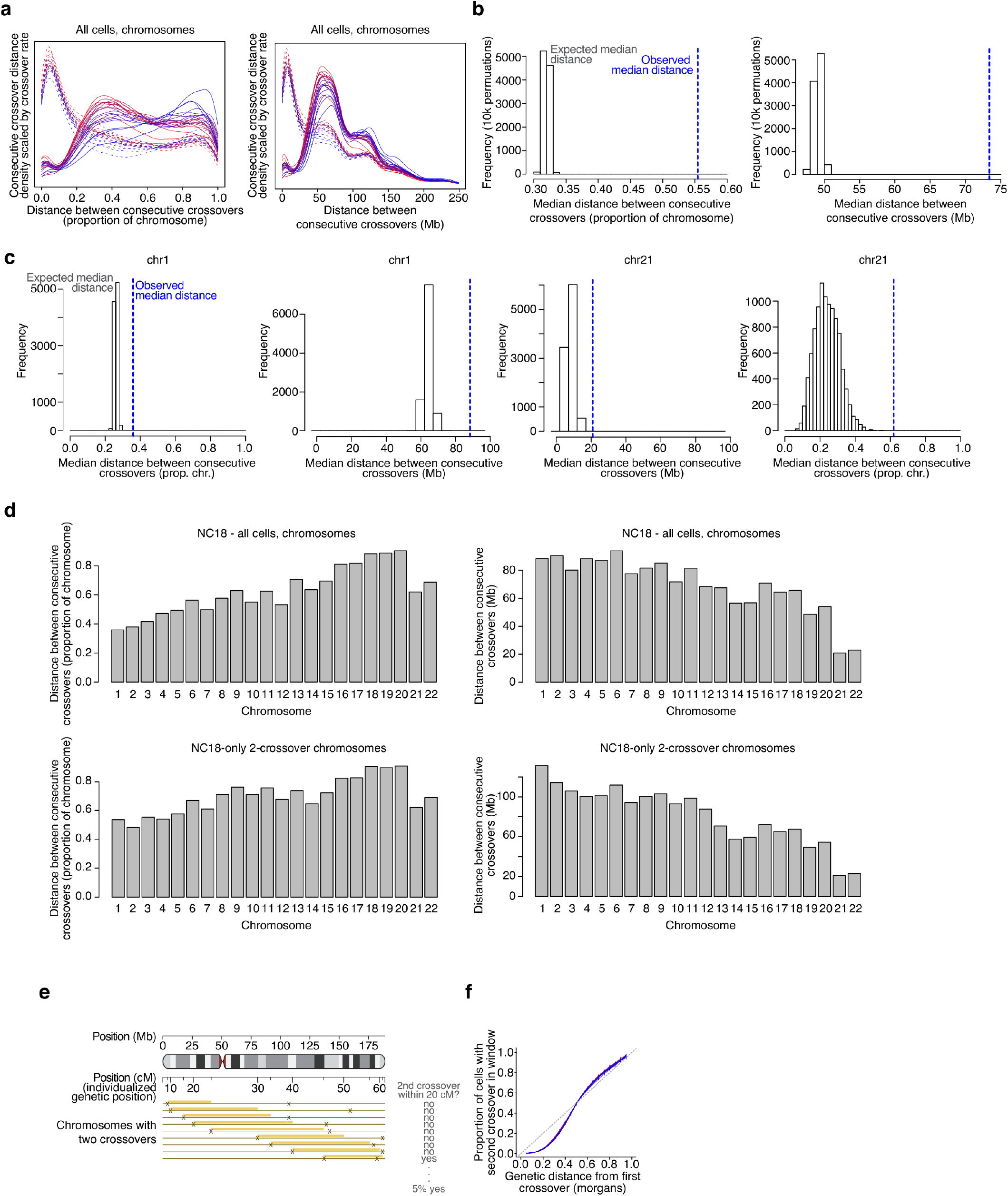
Crossover interference in individual sperm donors and on chromosomes. **a**, Solid lines show density plots (scaled by donor’s crossover rate) of the observed distance (separation) between consecutive crossovers as measured in the proportion of the chromosome separating them (left) and in genomic (Mb) distance (right), one line per donor. Dashed lines show the distance between consecutive crossovers when crossover locations are permuted randomly across cells to remove the effect of crossover interference. **b**, The median of observed distances between consecutive crossovers for one donor (NC18, 10^th^ lowest recombination rate of 20 donors; blue dashed line) is shown with a histogram of the medians of 10,000 among-cell crossover permutations (both permutation *p*s < 0.0001). Units, proportion of the chromosome (left) and genomic (Mb) distance (right). **c**, Crossover separation on example chromosomes; plots are as in (**b**). (Permutation *p* < 0.0001 for all chromosomes in all sperm donors except occasionally chromosome 21, where especially few double crossovers occur). **d**, Median distances between donor NC18’s consecutive crossovers for each autosome for all inter-crossover distances (top) and inter-crossover distances only from chromosomes with two crossovers (bottom). Units are proportion of the chromosome (left) and genomic (Mb) distance (right). **e**, Schematic: analyzing crossover interference in individualized genetic distance (one 20 cM window shown) using a donor’s own recombination map. **f**, When parameterized using each donor’s own genetic map, sperm donors’ crossover interference profiles across multiple genetic distance windows (as shown in **e**) do not differ (Kruskal–Wallis chi-squared = 0.22, *df* = 19, *p* = 1 using 20 estimates [cM distances] for each of 20 donors). Error bars, binomial 95% confidence intervals. This suggests that inter-individual variation in crossover interference, while substantial when measured in base pairs (as in **a**, **b**, **Fig. 4bd**, and Extended Data Fig. 12), is negligible when measured in genetic distance, pointing to a shared influence upon crossover interference and crossover rate.

**Extended Data Figure 12.**
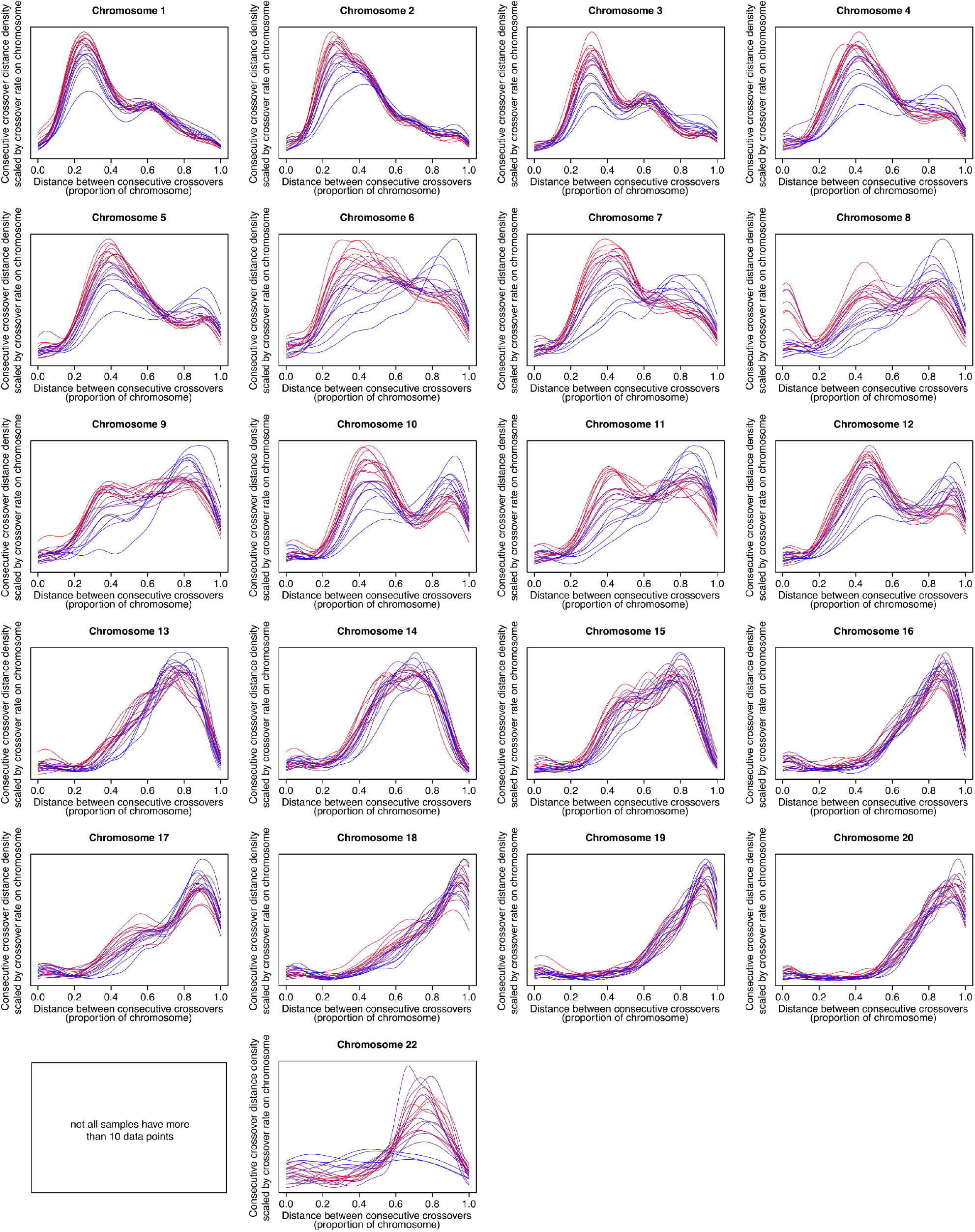
Per-chromosome crossover interference. (consecutive crossover separation density plots scaled by each donor’s crossover rate) for each of the 22 autosomes for each of the 20 sperm donors. All cells are included. Distance, proportion of the chromosome separating consecutive crossovers. (Extended Data Fig. 10b, left, shows all chromosomes combined; Extended Data Fig. 14 shows this with a different distance unit). Data is not shown for any chromosome(s) in which any donor had <10 chromosomes with ≤2 crossovers.

**Extended Data Figure 13.**
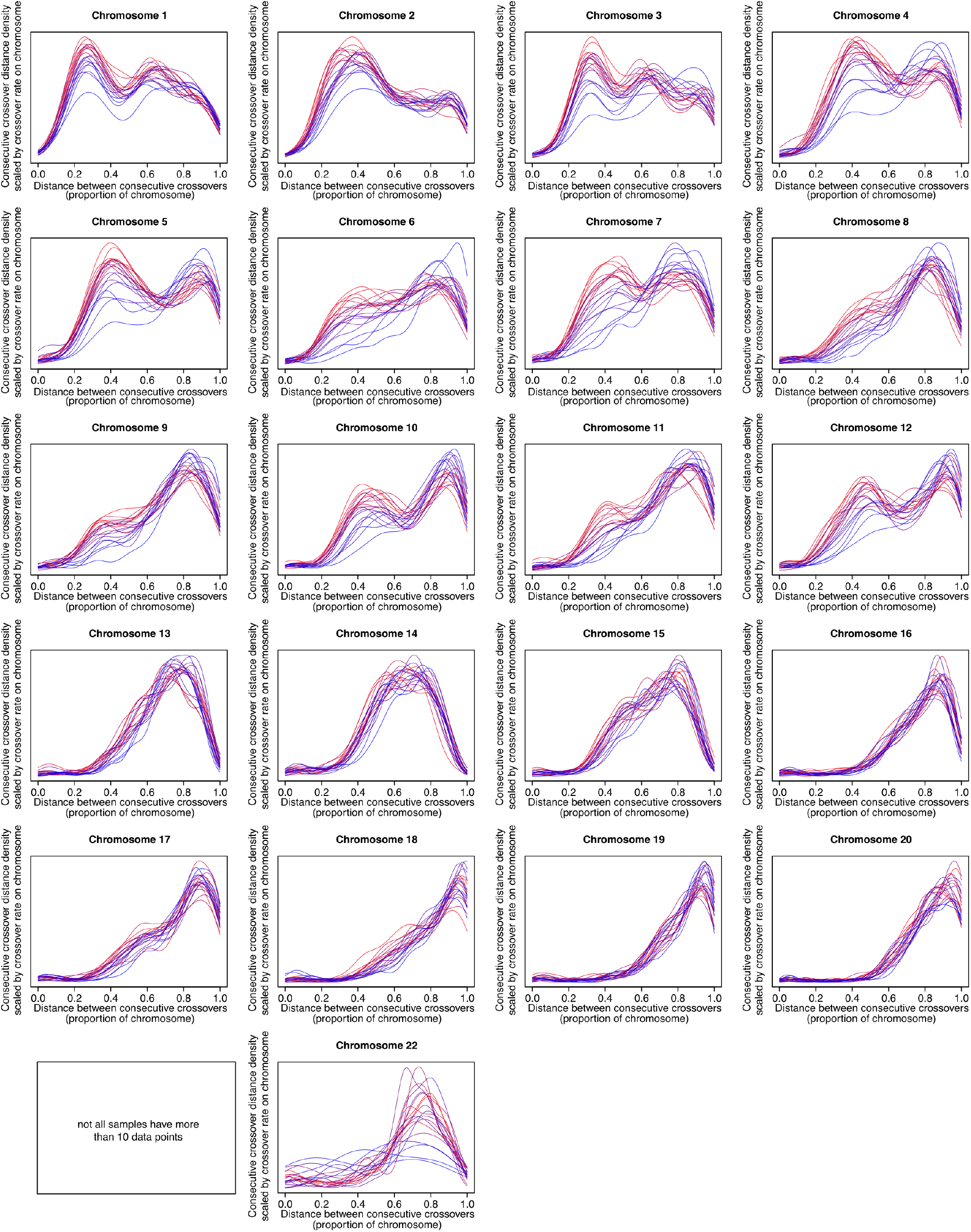
Per-chromosome crossover interference on two-crossover chromosomes. (consecutive crossover separation density plots scaled by each donor’s crossover rate) for each of the 22 autosomes for each of the 20 sperm donors. Only chromosomes with two crossovers are included. Distance, proportion of the chromosome separating consecutive crossovers (**Fig. 4b**, left, shows all chromosomes combined; Extended Data Fig. 15 shows this with a different distance unit). Data is not shown for any chromosome(s) in which any donor had <10 chromosomes with 2 crossovers

**Extended Data Figure 14.**
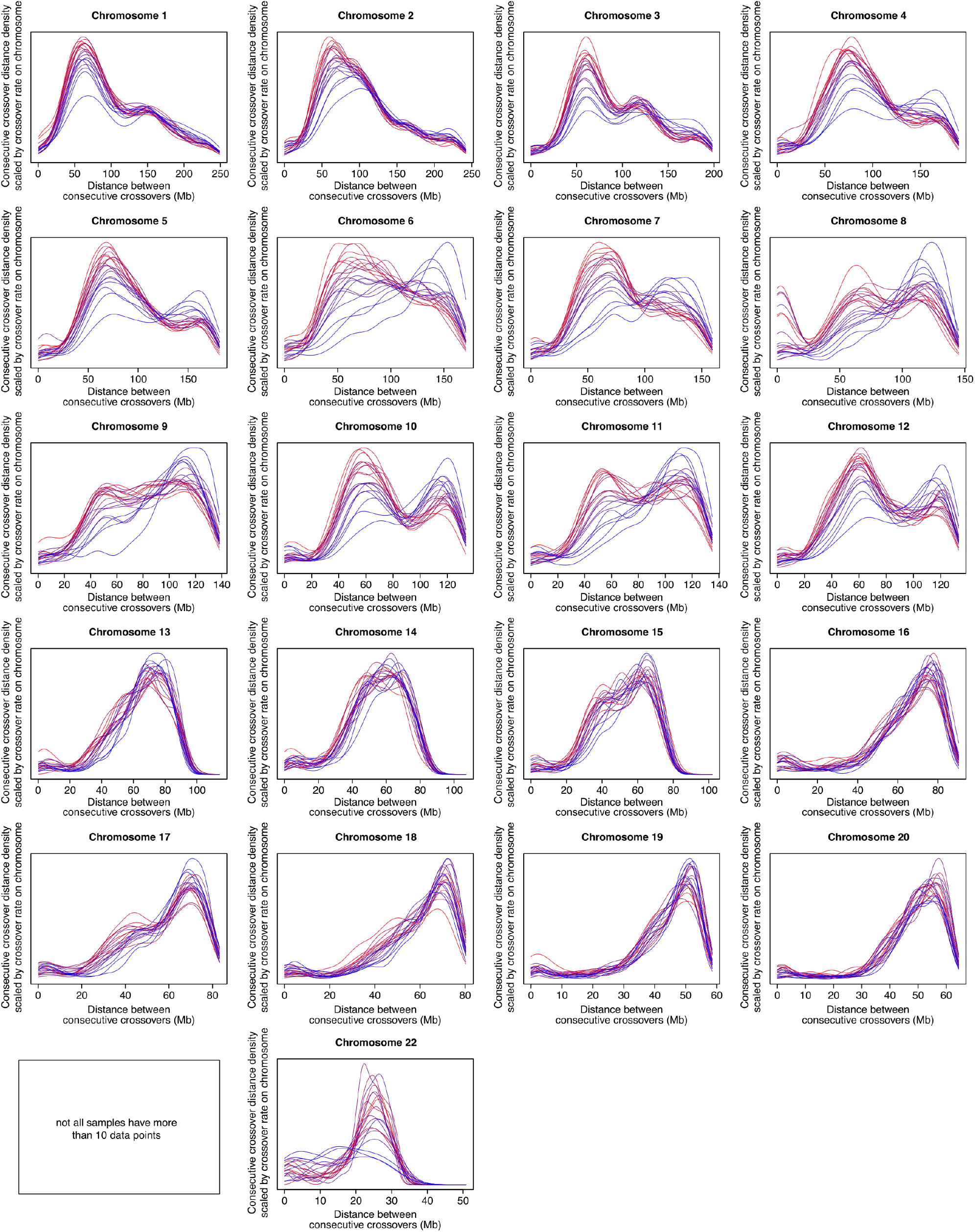
Per-chromosome crossover interference. (consecutive crossover separation density plots scaled by each donor’s crossover rate) for each of the 22 autosomes for each of the 20 sperm donors. All cells are included. Distance, genomic distance (Mb) (Extended Data Fig. 12 shows this with a different distance unit). Data is not shown for any chromosome(s) in which any donor had <10 chromosomes with ≤2 crossovers

**Extended Data Figure 15.**
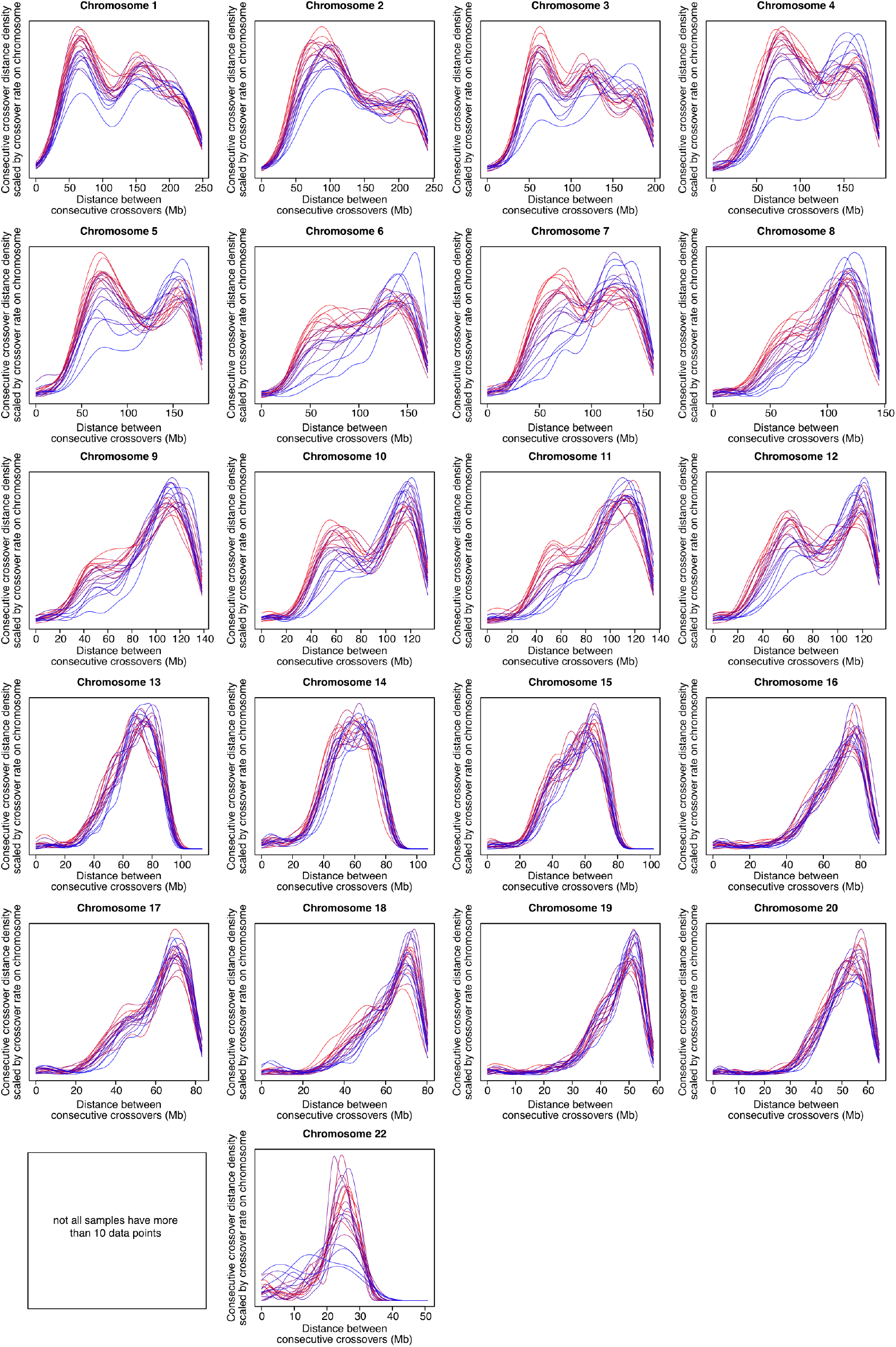
Per-chromosome crossover interference on two-crossover chromosomes. (consecutive crossover separation density plots scaled by each donor’s crossover rate) for each of the 22 autosomes for each of the 20 sperm donors. Only chromosomes with two crossovers are included. Distance, genomic distance (Mb) (Extended Data Fig. 13 shows this with a different distance unit). Data is not shown for any chromosome(s) in which any donor had <10 chromosomes with 2 crossovers

**Extended Data Figure 16.**
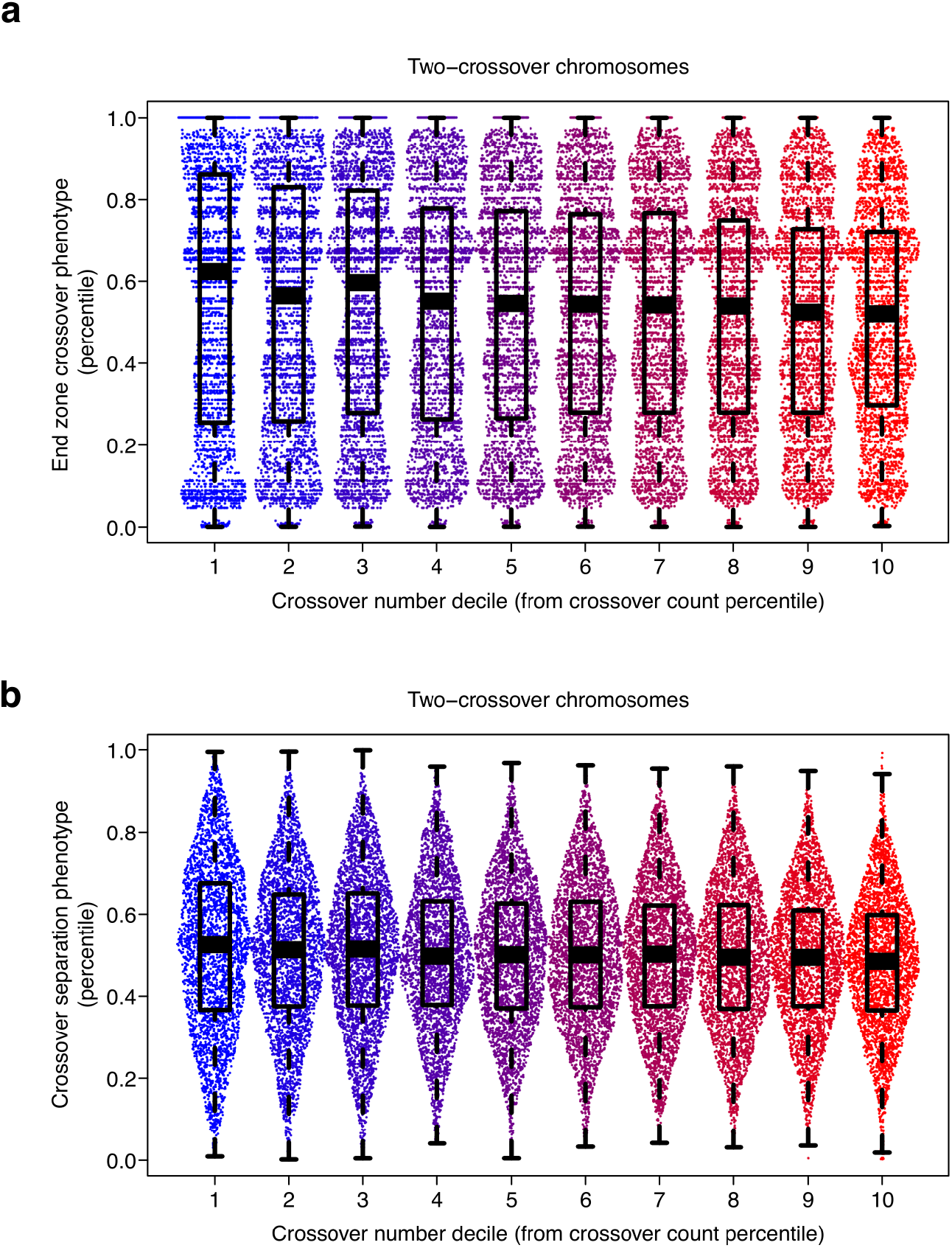
Crossover placement and interference on two-crossover chromosomes among cells with different crossover rates. Boxplots show medians and interquartile ranges with whiskers extending to 1.5 times the interquartile range from the box. Each point is a cell. **a**, Within-donor percentile of proportion of crossovers from two-crossover chromosomes falling in distal zones plotted vs. crossover rate decile. Data as in **Fig. 4d**, but showing all 10 deciles of crossover rate normalized within-sperm-donor by converting each cell’s crossover count to a percentile within-donor (All cells from all donors shown together, *n* cells in deciles 1-10: 3,152, 3,122, 3,276, 3,067, 3,080, 3,073, 3,135, 3,132, 3,090, 3,101 [31,228 total]). Because the initial data is proportions with small denominators (number of crossovers on all two-crossover chromosomes), an integer effect is evident as pileups at certain values. **b**, Crossover interference from two-crossover chromosomes (median consecutive crossover separation per cell shown). Data as in **Fig 4e** (each point represents the median of all percentile-expressed distances between crossovers from all two-crossover chromosomes in one cell, with percentile taken within-chromosome), groupings as in (**a**).

**Extended Data Figure 17.**
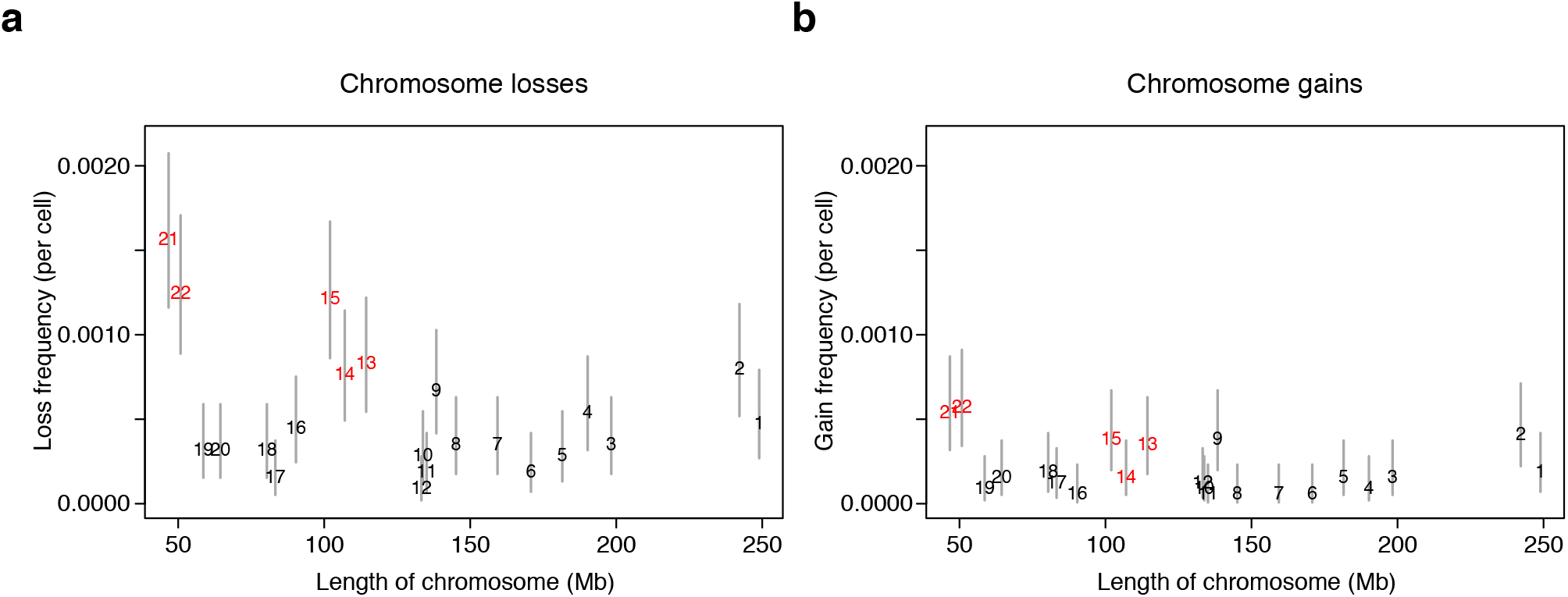
Aneuploidy frequency and chromosome size. The across-donor per-cell frequency of chromosome losses (left) and gains (right), as in **Fig. 5b**, plotted against the length of the chromosome (hg38; for losses, Pearson’s *r* = −0.29, *p* = 0.19 and for gains, Pearson’s *r* = −0.23, *p* = 0.30). Red labels, acrocentric chromosomes. Error bars, 95% binomial confidence intervals on per-cell frequency (number of events / number of cells, all 31,228 cells included).

**Extended Data Figure 18.**
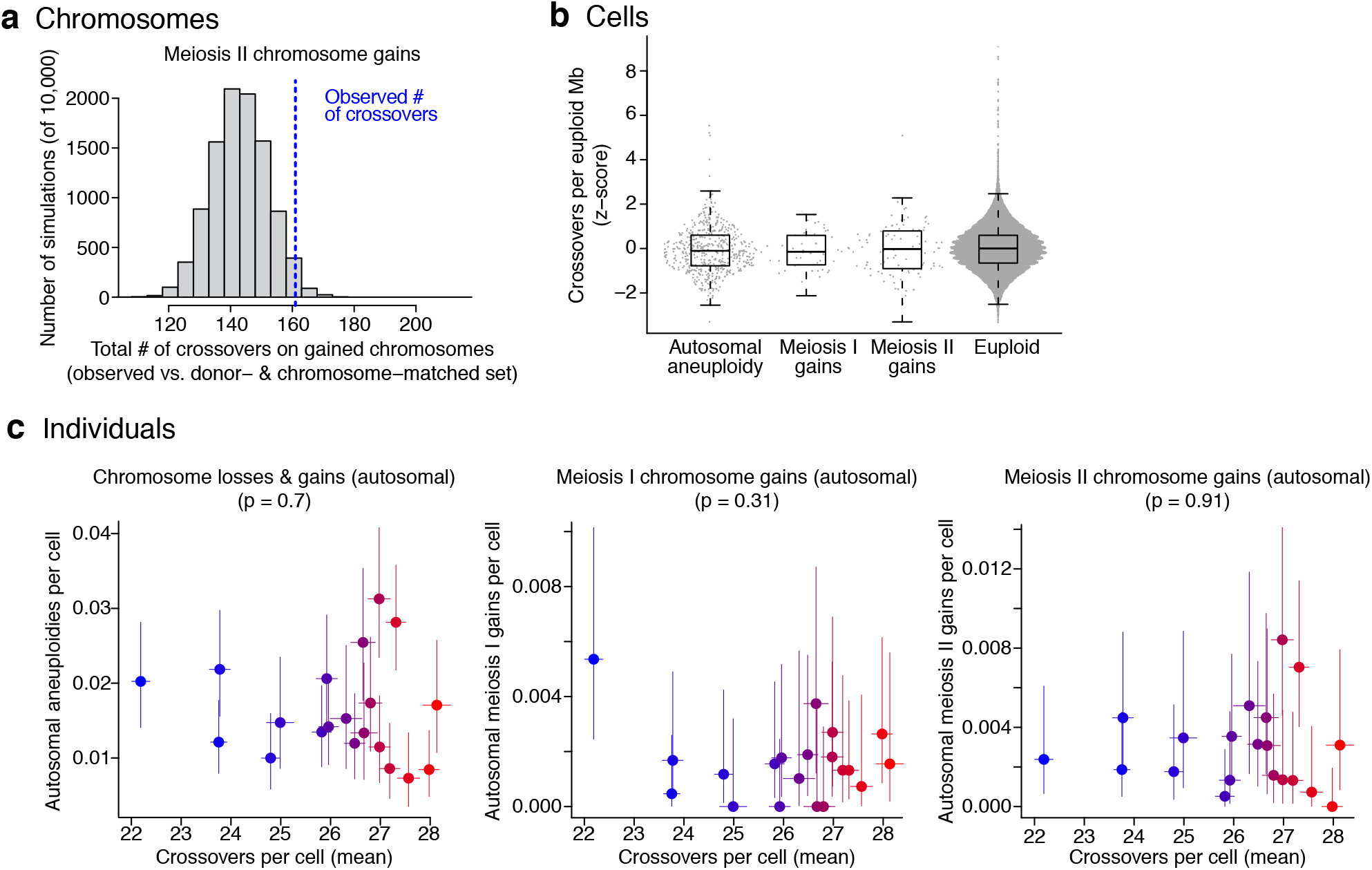
Relationship between crossover and aneuploidy frequencies across MII-gained chromosomes, cells, and donors. Only autosomal whole-chromosome aneuploidies are included. **a**, As in **Fig. 5f**, but for gains occurring during MII. Total inferred crossover number on gained chromosomes (blue line, summed across *n* = 71 MII-derived gained chromosomes of one whole copy from all individuals with fewer than 5 crossovers called on gained chromosome) compared to 10,000 donor and chromosome-matched sets (71 × 2 chromosomes per set) of properly segregated chromosomes (gray histogram). (One-sided simulation-derived *p* = 0.98 for MII, for the hypothesis that gained chromosomes have fewer crossovers; sister chromatids nondisjoined in MII capture all crossovers whereas matched chromosomes do not: matched simulations and homologs nondisjoined in MI capture only a random half of crossovers occurring on that chromosome in the parent spermatocyte). **b**, Crossovers per non-aneuploid megabase from each cell from each donor, split by aneuploidy status (*n* cells = 498, 50, 92, 30,609, left-to-right; “euploid” excludes cells with any autosomal whole- or partial-chromosomal loss or gain and “gains” includes gains of one or more than one chromosome copy; Mann–Whitney test *W* = 7,264,117, 722,191, 1,370,376; *p* = 0.07, 0.49, 0.66 for all autosomal aneuploidies, meiosis I (MI) gains, and meiosis II (MII) gains, respectively, all compared against euploid). Each cell is one point; boxplots show medians and interquartile ranges with whiskers extending to 1.5 times the interquartile range from the box. **c**, Per-cell crossover rates vs. per-cell aneuploidy (loss and gain) rates, with one point for each of the 20 donors (colored by crossover rate). *p* values shown in subtitles are for Pearson’s correlation tests. Error bars are 95% confidence intervals.

**Extended Data Figure 19.**
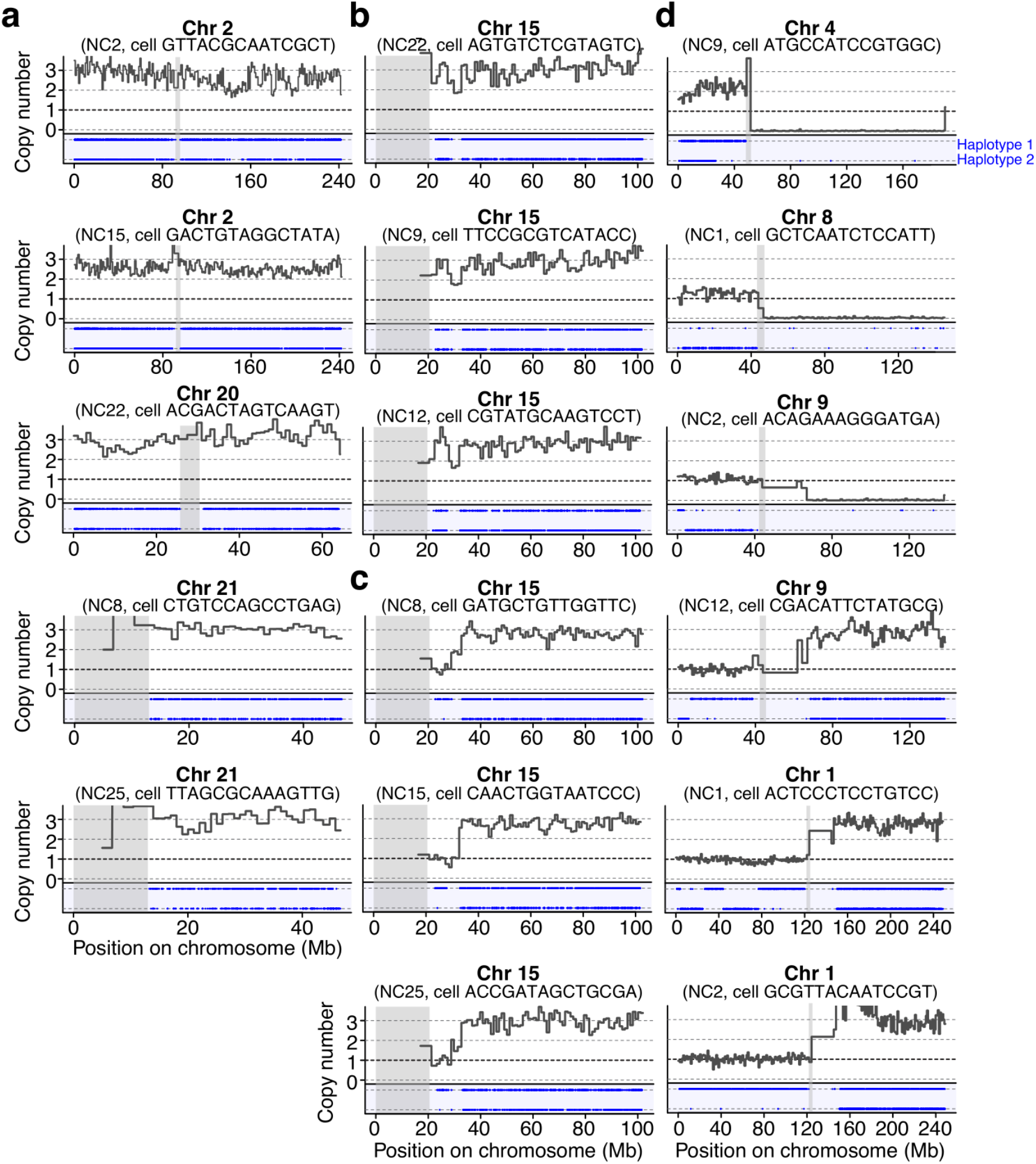
Further examples of non-canonical aneuploidy events detected with Sperm-seq, including those shown in Fig. 6. Copy number, SNPs, haplotypes, and centromeres are plotted as in **Fig. 5a**. Donor and cell identity are noted as subtitles. Coordinates are in hg38. Chromosomes 2, 20, 21 (**a**) and 15 (**b**) are sometimes present in an otherwise haploid sperm cell in 3 copies. **c**, A distinct triplication of chromosome 15, from ∼33 Mb onwards, but not including the first part of the *q* arm, also occurs in cells from 3 donors. **d**, Chromosome arm-level losses (top) and gains (including in more than one copy, bottom three panels, and a compound gain of the *p* arm and loss of the *q* arm, top panel).

