## Supplemental Text and Methods for "Insights about variation in meiosis from 31,228 human sperm genomes"

**for:**

#### Supplemental Text

##### Phasing validation and performance

Phasing of simulated single sperm data showed that phasing was 99.9% accurate when an average of 1% of heterozygous sites were covered in 1000 cells (Methods), similar to experimental coverage (**Table 1**). SNP coverage and the number of cells included affect phasing performance (**Extended Data Fig. 1a,b**). Comparison of experimental results to population-based phasing by Eagle[1,2] showed 97.5% phase concordance of consecutive heterozygous sites phased in both methods. Comparison to heterozygous SNP pairs in perfect linkage disequilibrium in population-matched 1000 Genomes[3] samples showed 97.9% concordance of experimental phase with linked alleles.

In this study, 97.3-99.98% (with a median across donors of 99.9%) of all called heterozygous sites were phased into chromosome length haplotypes; not all single SNPs were observed in enough cells to be phased.

##### Number and resolution of detected crossovers

Analysis of Sperm-seq data identified 813,122 crossovers in 31,228 gamete genomes (25,839–62,110 per sperm donor, **Table 1**). Previous human sperm cell sequencing and typing studies identified 2,000–2,400 crossovers[4,5], and the most recent single-sperm sequencing technology identified 24,672 crossovers in hybrid mice[6]. Recently, a very large pedigree-based study found 1,476,140 paternal crossovers in 56,321 paternal meioses[7].

The resolution of crossovers, which depends on density of SNP ascertainment in the cell and at the locus where they occur, was < 10 kb for 1.2% (9,746) of crossovers, < 100 kb for 23.0% (186,695), < 500 kb for 75.0% (610,121), < 1 Mb for 90.5% (735,955), and < 5 Mb for 99.7% (810,331).

**Quantification of similarity in genetic maps among sperm donors, HapMap, and deCODE**

Crossover rates (cM/physical distance) were correlated between sperm donors and between sperm donors and known genetic maps (pedigree-derived paternal map from deCODE[8] and population linkage disequilibrium-derived sex-averaged map from HapMap[9]). Among-sperm donor correlation (Pearson's *r*) in crossover rate (cM/physical distance) ranged from 0.62 to 0.88 at small 500 kb scale, and from 0.95 to 0.99 at larger 10 Mb scale, and the correlation between sperm donors and deCODE's recombination rates ranged from 0.66 to 0.86 at 500 kb scale and 0.92 to 0.96 at 10 Mb scale. Between sperm donors and HapMap's recombination rates, correlation coefficients ranged from 0.51 to 0.64 at 500 kb scale and 0.89 to 0.93 at 10 Mb scale. The individual genetic map from a previous single-sperm study[4] had similar correlation with population resources: in 3 Mb bins, Pearson's *r* correlation coefficients with HapMap sex-averaged and deCODE paternal maps were 0.71 and 0.77, respectively.

**Correlation of crossover number on different chromosomes in gametes**

Because crossover number is noisy within cells, a correlation of crossover number across chromosomes within cells could be hard to detect in our data. Moreover, in sperm cells, coordination of crossover number across chromosomes would occur in the primary spermatocytes undergoing meiosis, with its effects (crossovers) distributed randomly among the four daughter cells, resulting in a diffuse, hard-to-detect signal of small magnitude. To maximize power, we looked for this correlation between the number of crossovers in the largest possible equally sized sets of chromosomes (odd-numbered vs. even-numbered), recognizing that any observed correlation would likely substantially underestimate the biological effect size. Furthermore, we aggregated all 31,228 cells across all 20 donors by converting the total crossovers on all odd-numbered chromosome crossovers to a percentile and doing the same with the summed even-numbered chromosome crossovers. The correlation across these 31,228 cells was $r = 0.09$, $p = 8 \times 10^{-54}$.

All 20 individual donors had a positive Pearson's $r$ (sign test $p = 2 \times 10^{-6}$). Among donors (median $r = 0.1$, median $p = 3 \times 10^{-5}$), the donor with $r$ closest to median (NC4) had $r = 0.11$, $p = 3 \times 10^{-5}$; the donor with the smallest $r$ (NC12) had $r = 0.04$, $p = 0.09$; and the donor with the largest $r$ (NC10) had $r = 0.25$, $p < 10^{-14}$.

**The distribution of crossover number per cell vs. expected (random, independent) distribution**

The number of observed crossovers per chromosome per gamete exhibited less variance than expected relative to a purely random (Poisson) process, in which all crossovers are independent events. The observed median variance in crossover number across chromosomes and donors was 0.71, a 41% reduction relative to the median expected variance of 1.20 (this reduction was significant: one-sample chi-squared test on variance $p < 6 \times 10^{-10}$ for all donors and chromosomes). Additionally, fewer cells had chromosomes with no crossovers or many crossovers than would be predicted by a model in which crossovers are independent, random events (**Extended Data Fig. 7**) (all donors' and chromosomes' chi-squared test against the expected Poisson distribution $p < 2 \times 10^{-6}$; Methods).

**Control analyses for inter-individual differences in crossover interference and its correlation with crossover rate**

Different donors with different crossover rates had different chromosomal compositions of two-crossover chromosomes (*i.e.,* high–crossover rate donors may have few two-crossover chromosome 1s but many two-crossover chromosome 18s, whereas low crossover rate donors may have the reverse pattern). To determine whether the observation of individuals' crossover interference differences and the negative correlation between interference and crossover rate were robust to this compositional effect, we down-sampled each individual to have the same number of two-crossover chromosomes for each chromosome as the individual with the lowest number of two-crossover chromosomes for that chromosome (for example, NC26 had the minimum number of two-crossover chromosome 3s, 329, so for of all other donors, 329 two-crossover chromosome 3s were randomly chosen for the analysis). We performed this down-sampling five times, and in all cases, crossover interference still differed among individuals (Kruskal–Wallis test chi-square from 5,522 two-crossover chromosomes in each of the 20 donors, 1,158.3–1,231.1 [median, 1,175.5]; *p*-value, $2 \times 10^{-249} – 8 \times 10^{-234}$ [median, $2 \times 10^{-237}$]) and was still negatively correlated with crossover rate, with similar correlation coefficient as when all data were included (Pearson's *r* across 20 donors, -0.90 – -0.93 [median, -0.91]; *p*-value, $5 \times 10^{-9} – 6 \text{ x } 10^{-8}$ [median, $2 \times 10^{-8}$]).

In theory, the inter-individual difference in crossover interference difference and its negative correlation with crossover rate could be due to differential rates of failure to detect crossovers at the very ends of the chromosome, causing true three-crossover chromosomes to be included in the two-crossover chromosome pool. If this were to happen in a biased fashion (more often in higher recombination rate sperm donors), it could inflate the observed difference. To control for this possibility, we preferentially removed chromosomes with the shortest inter-crossover distances from the highest–crossover rate individuals (Methods); in this analysis, the inter-individual differences in crossover interference (Kruskal–Wallis chi-squared = 992, $df = 19$, $p = 3 \times 10^{-198}$, from *n* two-crossover chromosomes retained per donor = NC1: 5,337, NC10: 6,120, NC11: 104,57, NC12: 11,107, NC13: 8,450, NC14: 7,344, NC15: 9,171, NC16: 9,214, NC17: 8,186, NC18: 8,831, NC2: 8,268, NC22: 9,166, NC25: 12,392, NC26: 5,300, NC27: 7,019, NC3: 7,084, NC4: 8,084, NC6: 7,466, NC8: 9,144, NC9: 10,359) and negative correlation with crossover rate across 20 donors (Pearson's $r = -0.80$, $p = 3 \times 10^{-5}$) persisted.

**Inter-individual aneuploidy frequency variance**

The observed 4.5-fold variation in aneuploidy frequency across sperm donors could possibly derive from differences in statistical sampling. To investigate this possibility, we simulated the presence or absence of aneuploidy in each cell of each donor by drawing from the Poisson distribution with lambda equal to the total number of whole aneuploidies observed divided by the total number of cells observed across donors (787/31,228); each donor's simulation had the same number of cells as ascertained in that donor as in **Table 1**. For each simulation, we calculated the variance and median absolute deviation (MAD) across 20 simulated donors' aneuploidy frequencies; this process was repeated for a total of 10,000 simulations. We performed the same calculation for whole-chromosome losses and gains (lambda = 554/31,228 and 233/31,228, respectively). All observed across-donor variances and MADs were larger than the mean of the simulated variances (ratio of observed vs. simulated mean for variance: 4.5, 3.0, and 2.7 for all aneuploidies, losses, and gains, respectively; for

MAD: 1.7, 1.5, and 1.7), and the permutation tests were significant (variance: $p < 1 \times 10^{-4}$, $p < 1 \times 10^{-4}$, and $p = 2 \times 10^{-4}$ for all aneuploidies, losses, and gains, respectively; MAD: $p = 0.006$, $p = 0.037$, and $p = 0.007$).

**Is the observed excess of chromosome loss vs. gains likely to reflect a technical artifact?**

The observed overabundance of chromosome losses (or dearth of chromosome gains) could in theory be a technical artifact. If we over-detected losses, cells would most likely have lost chromosomes during sperm or droplet preparation. If so, this should be most common among short chromosomes, which might more easily become disentangled from the rest of the nucleus than long chromosomes. If we missed gains, cells containing them might have been excluded as cell doublets due to the presence of two haplotypes along a chromosome; consequently, gains of longer chromosomes, which contribute more to the global proportion of the genome containing two haplotypes, would be under-called. (To explicitly correct for this possibility, we removed the chromosome with the highest prevalence of both parental haplotypes from cell doublet.) Alternatively, heavier cells (with gains) might somehow be excluded from analysis, although we cannot currently explain why this might have occurred; if so, we would expect to see fewer gains of large chromosomes. However, none of these cases seem likely: chromosome length was not correlated with loss or gain frequency (for losses, Pearson's $r = -0.29$, $p = 0.19$ and for gains, Pearson's $r = -0.23$, $p = 0.30$, **Extended Data Fig. 17**). Additionally, we observed more losses than gains both on the sex chromosomes and on the autosomes, and sex chromosomes were not included in the doublet removal algorithm.

**Relationship between aneuploidy and crossover in chromosomes, cells, and individuals**

Because crossover calling on gained chromosomes is difficult (Methods), an excess of crossovers was sometimes called on individual gained chromosomes. We calculated the total number of crossovers both on 1) all gained chromosomes from MI ($n = 37$) or MII ($n = 87$) and 2) gained chromosomes with fewer than 5 crossovers called (from MI, $n = 32$, and MII, $n = 71$). We compared these totals to the total crossovers called in each of 10,000 sets of crossovers matched for chromosome and donor (and exclusion based on crossover number), where two chromosomes so matched were randomly chosen for each gain and all gains were included for one set. In both comparisons, MI gains had fewer total crossovers than matched sets (one-sided permutation $p = 0.0001$ for all gains, $p < 0.0001$ for gains with under 5 crossovers) and MII gains did not have fewer crossovers in total than matched sets (one-sided permutation $p = 1$ for all gains, $p = 0.98$ for gains with fewer than 5 crossovers).

The observed near-excess of crossovers on chromosomes gained in MII vs. matched sets occurs because the gain approximation (matching) is less appropriate for MII gains than MI gains. Sister chromatids fail to disjoin in MII gains, resulting in the presence of both sister chromatids of one homologous chromosome. These sister chromatids retain every crossover that happened on the parent chromosome in the parent spermatocyte, whereas chromatids from different homologs (like those gained in MI or in randomly chosen pairs of chromosomes) report on average half of the crossovers that happened on the parent chromosome. That is, MII gains report all physical crossover events (chiasmata) whereas non-sister chromatids report only chiasmata in which they were involved.

If factors that promote crossovers are generally protective against aneuploidy, individuals and cells with higher recombination rates would have lower aneuploidy rates. At the cell level, euploid and aneuploid gametes exhibited no differences in crossover frequency, nor did gametes with MI-derived or MII-derived chromosome gains (**Extended Data Fig. 18b**, Mann–Whitney test of crossovers per non-aneuploid megabase $W = 7{,}264{,}117$, 722,191, 1,370,376; $p = 0.07$, 0.49, 0.66 for all cells with whole-chromosome aneuploidy, MI whole-chromosome gains, and MII whole-chromosome gains vs. euploid, respectively; Methods). In addition, linear regression using aneuploidy status to predict crossover number in individual cells found no strong relationship between crossover rate and the rates of aneuploidy from either meiotic division (all aneuploidies $p = 0.33$, MI gains $p = 0.05$, MII gains $p = 0.26$; Methods). If the within-cell effect were of the magnitude of missing an entire chromosome's crossover complement from the non-aneuploid chromosomes in aneuploid cells, we would have been able to detect it: when we included aneuploid chromosomes (which obligately have 0 crossovers in our data unless specifically investigating gained chromosomes) in the analysis, we obtained significance in both the Mann–Whitney test and linear regression (all $p < 0.01$). Presumably, cells with aneuploidy occurring in MI would on average have slightly fewer total crossovers than euploid cells due to the observed slight correlation of crossover number across chromosomes.

Although the 20 individuals exhibited a 4.5-fold variation in aneuploidy rates and a 1.3-fold variation in crossover rates, these rates were not correlated with each other (Pearson's $r$ = -

0.09, $p = 0.70$) (**Extended Data Fig. 18c**, left). These rates remained uncorrelated when we focused on chromosome nondisjunctions occurring in MI (when crossovers occur) (MI: Pearson's $r = -0.24$, $p = 0.31$; MII: Pearson's $r = 0.03$, $p = 0.91$; **Extended Data Fig. 18c**, center and right). With 20 donors, we were 80% powered to detect an $r$ of 0.58 at $p = 0.05$.

**An excess of gains of more than one copy of chromosome 15**

More cells had three copies of chromosome 15 (potential double nondisjunction or unexplained events) than two copies of chromosome 15 (single nondisjunction events). Six cells had whole-chromosome triplications (as in **Fig. 6b, Extended Data Fig. 19b**), four cells had all of the *q* arm except for the pericentric region present in three copies (as in **Fig. 6c, Extended Data Fig. 19c**), and only two cells had gains of just one copy of chromosome 15. Twenty-two one-copy gains and no two-copy gains were expected from the Poisson distribution (total expected number of gains: sum of gained copies of chromosome 15 [22, 1 × 2 gains of one copy + 2 × 10 gains of two copies]; and total number of events: number of cells [31,228]), significantly different from our observations (Fisher's exact test $p = 2 \times 10^{-7}$).

#### Methods

*Our scripts are available via Zenodo, http://dx.doi.org/10.5281/zenodo.2581596. Scripts are referenced by name in the sections describing analyses they perform. Other tools are available as referenced.*

*All statistical analyses were performed in R unless otherwise noted.*

*All p-values reported in the main or supplemental text are two-sided unless otherwise noted.*

##### Sample information

Sperm samples from 20 anonymous, karyotypically normal sperm donors were obtained from New England Cryogenic Center under a Not Human Subjects determination from the Harvard Faculty of Medicine Office of Human Research Administration (protocol numbers M23743-101 and IRB16-0834). Donors consented at the time of initial donation for samples to be used for research purposes. (Specimens can be obtained from New England Cryogenic Center upon IRB approval.) Samples arrived in liquid nitrogen in "egg yolk buffer" or "standard buffer with glycerol" (no further buffer information provided) and were aliquoted and stored in liquid nitrogen in the same buffers until library preparation.

We do not know the precise age of these sperm donors because it is against sperm bank policy to release this information. However, per sperm bank policy, all donors are over 18 years old and younger than 38 years old at the time of donation. A different cohort would be required for analysis of any age effects.

All donor identifiers used in the paper were new identifiers created specifically for this study and are not linked to any New England Cryogenic Center identifiers.

##### Sperm cell library generation

*Sperm preparation: nuclei decondensation*

Sperm cells were washed and their nuclei decondensed to create accessible sperm nuclei "florets" using a combination of published decondensation protocols[10,11] with some modifications. Sperm aliquots containing >200,000 cells were thawed on ice and then washed by spinning for 10 minutes at 400 g at 4°C. After removal of the supernatant, the pellet was resuspended in 10 μL phosphate-buffered saline (PBS, Gibco/LifeTechnologies) and recentrifuged under the same conditions. After removal of the supernatant, the sperm pellet was resuspended in 2.5 μL of a sucrose buffer containing 250 mM sucrose (Sigma), 5 mM $MgCl_2$ (Sigma), and 10 mM Tris HCl (pH 7.5, Thermo Scientific). Tubes containing sperm aliquots were submerged in liquid nitrogen and immediately quick-thawed by holding them in a warm fist; a total of three freeze-thaw cycles were performed.

The freeze-thawed sperm solution was then combined with 22.5 μL decondensation buffer consisting of 113 mM KCl (Sigma), 12.5 mM $KH_2PO_4$ (Sigma), 2.5 mM $Na_2HPO_4$ (Sigma), 2.5 mM $MgCl_2$ (Sigma), and 20 mM Tris (Thermo Scientific) freshly supplemented with 150 μM heparin (sodium salt from porcine, Sigma H3393) and 1 mM beta-mercaptoethanol (BME, Sigma). The reaction was incubated at 37°C for 45 minutes. To allow enzymatic DNA amplification, heparin was inactivated by mixing the sperm solution with 0.5 U heparinase I (Sigma H2519) by gently pipetting and incubating at room temperature for 2 hours[12].

The sperm solution was moved to ice, and sperm floret concentration was determined by diluting 1:100 with PBS and staining with 1X SYBR I (Thermo Scientific), and then loading onto a hemocytometer and counting using the green fluorescence channel at 10x magnification.

*Single-sperm library preparation*

Droplets were prepared using the following modifications to 10X Genomics' GemCode (version 1[13]) User Guide Revision C (in place of steps 5.1–5.3.9); all reagents come from the 10X Genomics GemCode kit. Sperm were prepared for use as input by combining 10,833 sperm with ultrapure water to a final volume of 5 μL; 10,000 sperm were used for library generation. To each sperm sample was added 60 μL of a master mix containing 32.5 μL GemCode reagent mix, 1.5 μL primer release agent, 9.2 μL GemCode polymerase, and 16.8 μL ultrapure water, and then the same was mixed by gentle pipetting with a wide-bore multichannel pipette.

GemCode beads were prepared by vortexing at full speed for 25 seconds, and then diluted 1:11 with ultrapure water to a total volume of at least 90 μL of 1:11-diluted beads per sample. Per 10X's protocol, 60 μL of sample-master mix combination was added to the droplet generation chip, followed by 85 μL of freshly pipette-mixed 1:11-diluted bead mixture and 150 μL of fresh droplet generation oil.

Droplets were then generated and processed through library generation following 10X Genomics' GemCode (version 1) User Guide Revision C (starting with step 5.3.10 and continuing with the rest of section 5 and all of section 6).

**Sequencing**

Two libraries were generated for each of the 20 sperm donors, with eight total libraries generated in a batch (eight wells per 10X Genomics GemCode chip). Additional libraries were generated for four initial samples with low output (analyzed) cell counts. Libraries were sequenced on S2 200 cycle flow cells on an Illumina NovaSeq, with four or five libraries sequenced per flow cell. The read structure was 178 cycles read 1, 8 cycles read 2 (index read one), 14 cycles read 3 (index read two containing the cell barcode), and 5 cycles read 4 (unused; included to fulfill the NovaSeq's paired-end requirement).

**Bulk sequence data processing**

To convert the data to mapped BAM files with cell and molecular barcodes encoded as read tags, we used Picard Tools (http://broadinstitute.github.io/picard) and Drop-seq Tools (https://github.com/broadinstitute/Drop-seq/releases; see https://github.com/broadinstitute/Drop-seq/blob/master/doc/Drop-seq_Alignment_Cookbook.pdf for details on running many of the tools)[14]:

Illumina BCL files were converted to unmapped BAM files using Picard's ExtractIlluminaBarcodes and IlluminaBasecallsToSam with read structure 178T8B14T (cell barcodes, present in the i5 index, were incorporated as read 2 for ease of downstream processing). 10X Genomics GemCode index barcode sequences (four per sample) were supplied for de-multiplexing.

Next, BAMs were processed to include unique molecular identifiers (UMIs) and cell barcodes as read tags, and to exclude reads with poor-quality cell barcodes or UMIs; consequently, each read was retained as single-end with its 14-bp cell barcode stored in tag XC and its 10-bp molecular barcode/unique molecular identifier (UMI) stored in tag XM. We used the first 10 bp of read 1 as the UMI, as this sequence contains the random primer used to prime the read as well as the bases in the genome directly following the site where the primer bound. First, DropSeq Tools' TagBamWithReadSequenceExtended was called with BASE_RANGE=1-

14, BASE_QUALITY=10, BARCODED_READ=2, DISCARD_READ=true, TAG_NAME=XC, NUM_BASES_BELOW_QUALITY=1 to convert the second read (cell barcode) to tag XC and drop this read from the output BAM file, tagging reads with at least one cell barcode base having quality below 10 with tag XQ. Subsequently, TagBamWithReadSequenceExtended was called again with BASE_RANGE=1-10, BASE_QUALITY=10, HARD_CLIP_BASES=true, BARCODED_READ=1, DISCARD_READ=false, TAG_NAME=XM, NUM_BASES_BELOW_QUALITY=1 to convert the first 10 bases of read 1 (template read) into the molecular barcode, tagged XM, tagging reads with more than one molecular barcode base having quality below 10 with tag XQ. Finally, DropSeq Tools' FilterBAM was called with parameter TAG_REJECT=XQ to exclude reads with low-quality bases in either the cell or molecular barcodes.

Reads were aligned to hg38 using bwa mem[15]. BAMs were converted to FastQ using Picard's SamToFastQ, FastQ reads were aligned using bwa mem -M, and then unmapped BAMs were merged with mapped BAMs using Picard's MergeBamAlignment, with non-default options INCLUDE_SECONDARY_ALIGNMENTS=false and PAIRED_RUN=false. Reads were considered PCR duplicates if they had the same cell and molecular barcodes and mapped to the same start position as a higher-quality read (best quality read retained) using Drop-seq Tools' SpermSeqMarkDuplicates (part of Drop-seq tools v2.2 and above) with options STRATEGY=READ_POSITION, CELL_BARCODE_TAG=XC, MOLECULAR_BARCODE_TAG=XM, NUM_BARCODES=20000, CREATE_INDEX=true. (BAM file for all lanes and index sequences from the same sample were merged using Picard's MergeSamFiles prior to alignment and/or during duplicate marking with all BAMs given as input to SpermSeqMarkDuplicates.)

**Sperm donor variant calling and individual sperm cell genotyping**

For each donor, we pooled all reads from all libraries, including reads that did not derive from a barcode associated with a complete sperm cell (sperm cells had many reads per cell barcode; some barcodes were only associated with a few reads, which nonetheless derive from donor DNA, rather than with an entire haploid genome). Using GATK v3.7[16,17] in hg38, we followed GATK's best practices documentation for base quality score recalibration, gVCF generation using HaplotypeCaller (in DISCOVERY mode with -stand_call_conf 20), and joint genotyping with GenotypeGVCFs. We filtered variants by first selecting only SNPs for downstream use with SelectVariants -selectType SNP and flagging those with a QD score < 3 for exclusion using VariantFiltration (--filterExpression “QD<3.0”). We then performed VQSR following GATK’s best practices, except that we excluded annotations MQ and DP (VariantRecalibrator with GATK provided resources; -an QD, MQRankSum, ReadPosRankSum, FS, and SOR; -mode SNP; --trustAllPolymorphic; and tranches 90, 99.0, 99.5, 99.9, and 100.0). We applied tranche 99.9 recalibration using ApplyRecalibration -mode SNP and obtained the names of SNPs from dbSNP 146[18] using VariantAnnotator --dbsnp. Because we observed false positives even at lower tranches, we further filtered our sites to contain only biallelic SNPs present in Hardy–Weinberg equilibrium in 1000 Genomes Phase 3[3] using SelectVariants --concordance with a VCF containing only these sites (from GATK’s resource bundle, which contained a lifted-over VCF when used). To narrow to the final set of heterozygous SNPs used in phasing and single-sperm analysis, we also excluded SNPs in centromeric regions or acrocentric arms as defined by the UCSC Genome Browser’s cytoband track[19,20] (http://genome.ucsc.edu), and those in known paralogous regions as lifted over from Genovese et al 2014[21] (available upon request), and selected only heterozygous SNPs using SelectVariants -selectType SNP --selectTypeToExclude INDEL --restrictAllelesTo BIALLELIC --excludeFiltered --setFilteredGtToNocall --selectexpressions 'vc.getGenotype("""<sample name>""").isHet()'.

We identified which of these SNPs were present in each sperm cell and which allele was present using the tool GenotypeSperm (part of Drop-seq tools v2.2 and above, https://github.com/broadinstitute/Drop-seq). First, we generated an interval file for each heterozygous SNP in the donor’s genome using Drop-seq Tools’ CreateSNPIntervalFromVCF. We determined the number of reads and molecular barcodes covering each base at each heterozygous SNP using GenotypeSperm with INTERVALS= the previously-generated interval file for the donor and cell barcodes (CELL_BC_FILE) expected to correspond to full haploid genomes (identification described in the next section).

For downstream analyses (identifying doublets, crossover calling), we generated a file with columns cell, pos, and gt, with gt having the value 0 for the reference allele and 1 for the alt allele by including SNPs that had one or more UMIs covering only one base (from GenotypeSperm) matching the reference or alternate allele (from GATK). (See our script *gtypesperm2cellsbyrow.R.*)

#### Generation and validation of chromosome-length phased haplotypes

To phase sperm donors' genomes, we used all quality-controlled sites (as described above) from all cell barcodes expected to correspond to sperm cells. We identified barcodes potentially associated with cells by plotting the cumulative fraction of reads associated with each ranked barcode and identifying the inflection point of this curve, i.e., the point at which including more barcodes only marginally increased the proportion of total reads included, such that each subsequent barcode was associated with few reads (we used DropSeq tool's BamTagHistogram to obtain ranked read counts per cell barcode). We further narrowed this set to include only barcodes with substantial read depth on either the X or the Y chromosome but not both, as the vast majority of sperm cells should contain only one sex chromosome; thus, most barcodes associated with both the X and the Y chromosome likely captured two or more sperm cells. (We later added all these barcodes back in before formally identifying doublet cell barcodes).

To phase, for each chromosome we converted per-cell SNP calls into "fragments" for input into the HapCUT phasing software[22,23] by considering each consecutive pair of SNPs observed in a cell to be a fragment (see our script *gtypesperm2fmf.R*). We then used HapCUT with parameter –maxiter 100 to generate phase blocks; all phase blocks generated are the length of entire chromosomes. After identifying and removing cell doublets (see below), we repeated phasing with only non-doublet cell barcodes to correct any possible phase errors resulting from the inclusion of cell doublets.

We validated our phasing method in several ways. First, we simulated single-cell SNP observations from known haplotypes, including 2% genotype errors and a variable percentage of cell doublets (**Extended Data Fig. 1a,b**). Briefly, sites were randomly sampled from one known haplotype of chromosome 17 until a crossover location probabilistically assigned based on the deCODE recombination map[8], then randomly sampled from the other haplotype (for simplicity, one crossover was simulated per cell, consistent with crossover expectation on a short chromosome). To simulate PCR or sequencing errors, after the entire chromosome was simulated, 2% of the sites were randomly assigned to an allele. Doublets were simulated by combining two cells and retaining 70% of the observed sites at random. (See our script *simulatespermseqfromhaps.py*.) We performed five random simulations for each doublet proportion, mean proportion of total chromosomal sites "observed" in each cell, and total number of cells simulated, and then followed our phasing protocol on the simulated cell sets. We calculated the proportion of sites with the same allele from phasing with our simulated cells as the input haplotypes (unphased alleles counted as incorrect).

We also phased an initial sperm donor's genome from actual Sperm-seq data and compared these phased haplotypes to haplotypes generated for this donor using the population-based phasing algorithm Eagle[1,2]. We compared the phase relationship between each consecutive pair of SNPs (identifying the proportion of switch errors between the two phased sets). We further examined all pairs of alleles in perfect linkage disequilibrium in 1000 Genomes Phase 3[3] in the populations matching the donor's ancestry to determine whether these alleles occurred on the same experimentally derived haplotype.

**Identification of cell doublets**

To identify cell barcodes associated with more than one sperm cell (cell doublets), we identified the haplotype of origin for all observed autosomal SNPs in each cell barcode and then counted the number of times consecutively observed SNP alleles appeared on different parental haplotypes, which could occur because of crossover, error, or the presence of two haplotypes in the same droplet (doublet). We ranked barcodes by the proportion of consecutive SNPs that spanned haplotypes in this way using all SNPs from all autosomes except the autosome with the most haplotype-spanning consecutive SNPs (so as to avoid mistakenly identifying cells with chromosome gains as doublets); this resulted in a clear inflection point wherein cell doublets had a higher, quickly accelerating proportion of haplotype-spanning consecutive SNPs (**Extended Data Fig. 1a**). All cell barcodes below this inflection point (identified with the function *ede* from the R package *inflection* https://CRAN.R-project.org/package=inflection) were considered non-doublet (**Extended Data Fig. 1b**). (See our script *computeSwitchesandInflThresh.R.*)

**Identification of crossover events**

We identified crossover events on all autosomes, excluding the *p* arms of acrocentric chromosomes (as SNPs on these arms were excluded from analysis), by assigning each observed SNP in each non-doublet cell to its parental haplotype and finding transitions between these haplotypes using a Hidden Markov Model written in R with package *HMM* (https://CRAN.R-project.org/package=HMM) (**Fig. 2a**). To ensure that we detected crossovers located near the ends of SNP coverage (as sub-telomeric regions are known to be frequently used for crossovers in spermatogenesis), we ran the HMM both in the forward chromosomal and reverse-chromosomal directions, with start probability for one haplotype equal to 1 if the first two SNPs observed were of that haplotype. In addition to two states for parental haplotypes, we included a third "error" state to capture cases in which a haplotype 1 allele is observed in a haplotype 2 region (and vice versa), *e.g.*, due to PCR or sequencing error, gene conversion, or cases in which a small piece of off-haplotype ambient DNA was captured in a droplet. Crossovers were identified as regions where one haplotype transitions to another, or where one haplotype transitions into the error state and then into the other haplotype; crossover boundaries were defined as the last SNP in the first haplotype and the first in the next (with up to a few intervening "error" SNPs when boundaries were unclear). The key parameters for this algorithm are the transition probability between haplotypes (set to 0.001, from the per-cell median 26 crossovers divided by the per-cell median 24,710 heterozygous SNPs per cell per donor) and transition into and out of the "error" state (we set transition probability into this state to 0.03 from either haplotype, as only a few percent of SNPs are off-haplotype; we set the probability of staying in error to a higher value, 0.9, to allow for the occasional tract of SNPs from an ambient piece of off-haplotype DNA). Emission probabilities are 100% haplotype 1 alleles from haplotype 1, 100% haplotype 2 alleles from haplotype 2, and equal probability haplotype 1 or haplotype 2 alleles from the third "error" state. Crossover calling was robust with respect to transition probabilities so long as the transition probability remained low. (See our script *spseqHMMCOCaller_3state.R*, which calls crossovers on one chromosome.)

After aneuploidy identification, we marked aneuploid chromosomes as having no crossovers for all crossover analyses (absent chromosomes have no crossovers and crossover calling requires a special procedure on gained chromosomes, described in a later section.).

**Restricting to cell barcodes with coverage of the entire genome**

To examine evenness of coverage and enable aneuploidy identification, we used Genome STRiP (http://software.broadinstitute.org/software/genomestrip/) [24,25] to determine sequence read depth (observed number of reads divided by expected number of reads) in 100-kb uniquely mappable bins (which may be larger than 100 kb of chromosomal territory) across the genome in each sperm cell, using Genome STRiP's default GC bias correction and repetitive region masking for gr38. We divided this read depth by 2 to obtain read depth per haploid rather than diploid genome. Input to Genome STRiP was a BAM file containing only cells of interest with read groups set to <sample name>:<cell barcode> (created using Drop-seq Tools' ConvertTagToReadGroup with options CELL_BARCODE_TAG=XC, SAMPLE_NAME=<name of sample/donor>, CREATE_INDEX=true, and CELL_BC_FILE pointed to a list of cell barcodes potentially associated with cells, described above).

Although read depth was usually centered at 1 across chromosomes, we noted that a minority of cell barcodes were associated with eccentric read depth across many chromosomes, with read depth vacillating between 0 and 2 or more in waves. (We hypothesize that these cell barcodes were associated with sperm cells that did not properly decondense, such that some regions of the genome were more accessible than others, leading to undulating read depth due to transitions between more and less accessible chromatin.) To identify and exclude such barcodes, we treated read depths across each chromosome as a time series and used Box-Jenkins Autoregressive Integrated Moving Average (ARIMA) modelling (implemented via the R package *forecast*[26,27], excluding differencing) to model how read depth observations relied on their previous values (as in "wave" cases) and their overall averages. By visual inspection, we determined that chromosomes with certain ARIMA criteria were likely to be unstable in read depth, and that cell barcodes with five or more such identified chromosomes were likely to have eccentric read depth globally. We flagged individual chromosomes if 1) The sum of AR1 and AR2 coefficients was greater than 0.7, the AR1 coefficient was greater than 0.9, or the net sum of all AR and MA coefficients was greater than 1.25 and 2) either the net sum of AR and MA coefficients was greater than 0.4 or the intercept was less than 0.8 or greater than 1.2. If both criteria in (2) were met, this signified an exceedingly odd chromosome, which was therefore counted twice. Cell barcodes with five or more chromosomes flagged in this way were excluded from downstream analyses. (Because gains of large amounts of the genome cause artificially depressed read depths on non-gained chromosomes, we manually examined any cells with a large range of ARIMA intercepts and over five chromosomes denoted as unstable. Any such cells that had simply gained a large proportion of the genome, *e.g.*, 3 copies of chromosome 2, were included rather than excluded.) We initially cross-referenced all cell exclusions with called aneuploidies, confirming that cells were not excluded simply on the basis of having lost or gained a chromosome.

(See our scripts *setupgsreaddepth.R*, *exclbadreaddepth_arima_1.R, exclbadreaddepth_initid_2.R*, and *exclbadreaddepth_finalize_3.R*)

**Identification and use of replicate barcodes ("bead doublets")**

In addition to cases in which two sperm cells are in the same droplet with the same cell-barcoded bead (see "Identification of cell doublets"), it is possible for one sperm cell to be encapsulated in a droplet with more than one barcoded bead. We determined the proportion of SNPs that were of the same haplotype for each pair of barcodes to identify cases where pairs of sperm genomes were identical. Because barcodes capture different sets of SNPs, we imputed the haplotype of all heterozygous SNPs based on the haplotype of surrounding observed SNPs and locations of recombination events and compared all SNPs across sperm cell pairs. SNPs occurring between identified boundaries of crossovers were excluded from analysis. Sperm cells shared on average 50% of their genomes, with differential sharing normally distributed except for a few high-outlying sets of barcodes that shared nearly 100% of their SNP haplotypes (**Extended Data Fig. 2a**) – these were considered to represent "bead doublets" or replicate barcodes. All downstream analyses were performed on a unique set of cell barcodes, *i.e.*, only one barcode from a set corresponding to the same cell was chosen randomly and used for analysis. (See our scripts *imputeHaplotypeAllSNPs.R, compareSpermHapsPropSNPs.R, combineChrsSpermHapsPropSNPs.R*, and *curateNonRepBCList.R*)

We used these bead doublets to examine the reproducibility of SNP and crossover calling (**Extended Data Fig. 2c-e**). We determined the proportion of observed SNPs shared by replicate barcodes and of these, the proportion in which the same haplotype was detected; we then compared this to the same metrics for randomly chosen sets of cell barcodes. We also calculated how many of the crossovers observed in either of the two barcodes overlapped with crossovers in the other barcode, and for any non-overlapping crossovers, determined whether they occurred within 15 SNPs of the end of SNP coverage (suggesting random fluctuations at the end of coverage among barcodes), whether two crossovers were close to overlapping but were simply separated by one SNP, or whether they did not overlap for other reasons.

**Investigation of whether unequal SNP coverage impacts crossover analyses**

Because coverage of heterozygous sites is non-uniform across different cells from a given sperm donor and across sperm donors, it is possible that some inter-cell and inter-individual differences in crossovers could derive from this differential coverage and the resultant differential ability to call crossovers, or from different genomic resolution of called crossovers. To determine whether this affected our conclusions, we randomly downsampled SNP observations from each chromosome in each cell to have the same number of observed heterozygous sites, simply masking any number greater than this, and excluded cells with more than two chromosomes with fewer heterozygous sites. After this down-sampling, 98.6% of cells ($n$ = 30,778) were retained. We down-sampled to the number of SNPs per chromosome from the 25$^{th}$ smallest cell from NC26, the donor with the lowest median per-cell SNP observation count, for a total of 13,036 SNPs per cell. We chose the 25$^{th}$ smallest cell to avoid any potential systematic issues with the very smallest cells, while still retaining most cells. (See our scripts *getNSNPsPerChrForDownsample.R, downsampleCellsByRow.R,* and *getBCsWithEnoughChrs.R.*)

We then re-called crossovers from these SNPs ($n$ = 785,476), and determined the correlation between the number of crossovers (per cell and per chromosome) in these calls from equal SNP coverage and our initial, full-coverage calls (**Extended Data Fig. 3a,b,d**). We compared the locations of crossovers called from both SNP sets via Kruskal–Wallis tests comparing each chromosome's median position of crossovers (from all cells combined, with each chromosome's position distribution tested separately, all crossovers included) (**Extended Data Fig. 3c,e**). To directly confirm that the same conclusions were reached in analyses using both datasets (data not shown), we also performed most crossover analyses using crossovers called from both SNP sets.

**Crossover rate analyses**

*Comparison of crossover number distribution among cells to the Poisson distribution*

Based on the total number of crossovers observed across all cells for each sperm donor, we determined the expected number of cells with each crossover count if crossovers were distributed randomly among cells according to the Poisson distribution (lambda = total number of crossovers / total number of cells). For this purpose, we used the Poisson density function in R multiplied by the total number of cells to obtain counts with quantiles ($x$) spanning the minimum and maximum numbers of crossovers where the Poisson expectation rounded to be greater than 0 (we extended the analysis to the minimum [maximum] observed crossover count if this was lower [higher] than would otherwise be included). To directly compare the observed and expected (Poisson) distributions of crossovers per cell, we used a chi-squared test. We also determined the experimental (observed) and expected variance and kurtosis (variance of Poisson is lambda and kurtosis is 3+1/lambda; observed kurtosis was calculated with the kurtsosis function from the R package *moments* https://CRAN.R-project.org/package=moments). We tested whether the observed variances differed from the expected variances using a one-sample chi-squared test on variance as implemented with the function *varTest* in the package *EnvStats* (https://CRAN.R-project.org/package=EnvStats). We performed this analysis for each chromosome separately. (See our script *coRateVariationAnalysis_poisson.R*.)

*Correlation of crossover rate across gametes from the same donor*

To determine whether the (noisy) crossover rate correlated across chromosomes in sperm cells from the same donor, we looked for a correlation between the number of crossovers in the largest possible equally sized sets of chromosomes (odd-numbered vs. even-numbered) in each donor. We also aggregated across donors by converting each crossover sum (odd- and even-numbered chromosomes) to a percentile within each donor, and then combining all donors and performing a correlation test on these percentiles. (See our script *rateVOtherPtypesAcrossCellAggs.R*, which performs these and many other analyses.)

*Comparison of this study to population-based genetic maps*

To determine how our individualized genetic maps compared to genetic maps generated from population data, we obtained population genetic maps from HapMap[9] (sex-averaged) in hg38 from the Eagle phasing package[1,2] and from deCODE[8] (male-specific) in hg18. We converted the deCODE map to hg38 using UCSC Genome Browser's[19] Batch Coordinate Conversion (liftover), and dropped liftover failures, as the sequential nature of a genetic map means it is not damaged by missing SNP observations. Because we observed different heterozygous sites across sperm donors, we determined the genetic positions in 500-kb interval bins individually for each of the sperm donors. We determined the number of crossovers occurring before each 500-kb position, divided this number by the total number of sperm cells analyzed, and multiplied by 100 to get each 500-kb physical bin's location in centimorgans, thereby standardizing across donors. We found the genetic positions corresponding to these physical bins in HapMap and deCODE by identifying the closest typed SNP to each bin boundary, and then examined these standardized maps together (**Fig. 2d,e, Extended Data Fig. 6**). From these 500-kb genetic maps, we determined the recombination rate in intervals of various sizes for each donor, HapMap, and deCODE and correlated these rate profiles across samples. (See our scripts *computeGenDistsMultSamps.R* and *plotAnalyzeGenDists.R*)

**Identification and use of crossover zones**

To define territories of recombination use (**Extended Data Fig. 8**), we found local minima of the density (built-in function in R) of all crossovers' median positions across all samples on each chromosome. Minima were identified using the *findPeaks* function (from https://github.com/stas-g/findPeaks) on the inverse density with m=3. Crossover zones run from the beginning of the chromosome (including the whole *p* arm for acrocentric chromosomes) to the location of the first local minimum, from the location of the first local minimum plus one basepair to the next local minimum, and so on, with the last zone on each chromosome ending at the last basepair position of that chromosome. (See our script *findcozones_peaks.R*.)

To determine what proportion of crossovers occurred in the most distal (telomeric) zones, we divided zones into "end" and "not-end" groups; all zones that encompassed a telomere were defined as end zones. Acrocentric chromosomes have only one end zone because the *p* arm was excluded from analysis, whereas all other chromosomes have two end zones, and chromosomes with only one zone on the *p* arm and one on the *q* arm comprise only end zones. To obtain a per-cell proportion of the distal crossovers metric, we divided the total number of crossovers in each cell with midpoints in these end zones by the total number of crossovers in each cell (**Extended Data Fig. 10a**, left). To obtain a per-sperm donor proportion of crossovers in distal zones metric (as in **Extended Data Fig. 10a**, right), we divided the total number of crossovers across all cells with midpoints in the end zones by the total number of crossovers detected across all cells from that donor. To get comparable numbers when controlling for crossover rate by restricting analyses to chromosomes with two crossovers, for each cell (**Fig. 4d**), we divided the total number of crossovers from two-crossover chromosomes by the number of these crossovers that occurred in distal chromosomal zones. For each sperm donor (**Fig. 4b**), we divided the total number of crossovers in end zones from two-crossover chromosomes in any cell by the total number of crossovers from two-crossover chromosomes in any cell. (*n* two-crossover chromosomes included per donor = NC3: 7,848, NC9: 11,509, NC6: 8,234, NC25: 13,590, NC13: 9,280, NC4: 8,838, NC8: 9,952, NC27: 7,645, NC26: 5,741, NC14: 7,942, NC18: 9,509, NC1: 5,745, NC22: 9,816, NC17: 8,766, NC11: 11,104, NC10: 6,432, NC15: 9,618, NC16: 9,481, NC12: 11,420, NC2: 8,268.)

**Analysis of crossover interference across donors**

We looked for crossover interference in each donor by computing the distance between all consecutive pairs of crossovers on the same chromosome in the same cell (using the midpoint between the border SNPs as the position of each crossover). We expressed this distance both in base pairs and as the proportion of the non-centromeric chromosome (or non-acrocentric arm for acrocentric chromosomes) separating each consecutive crossover pair. To determine whether this distribution reflected crossover interference, we compared its median to the median distances between consecutive crossovers computed by permuting crossovers' cell identities 10,000 times (in a fashion similar to that used by Wang et al[5]). In this permutation, we randomly assigned crossovers to cells while keeping constant the distribution of the number of cells with each number of crossovers per cell (accomplished by permuting within-chromosome such that chromosome 1's distribution of chromosomes with 1, 2, 3… crossovers was maintained) and then computed each inter-crossover distance and the median of this distribution. We compared the observed median to the 10,000 permuted medians. We performed this process globally (combining all chromosomes and on each chromosome (**Extended Data Figs. 11b,c**). To determine whether the 20 samples differed in crossover interference, we used a Kruskal–Wallis test on all inter-crossover distances (**Extended Data Fig. 10b**; *n* inter-crossover distances per donor = NC3: 13,832, NC9: 20,125, NC6: 14,049, NC25: 22,918, NC13: 14,913, NC4: 14,516, NC8: 16,254, NC27: 12,200, NC26: 9,277, NC14: 12,795, NC18: 14,971, NC1: 9,165, NC22: 15,239, NC17: 13,515, NC11: 17,163, NC10: 9,499, NC15: 13,792, NC16: 13,134, NC12: 15,803, NC2: 10,519). We also performed these analyses on chromosomes with two crossovers (one inter-crossover distance per chromosome, *n* two-crossover chromosomes included per donor described above in "Identification and use of crossover zones") (**Fig. 4c**). (See our scripts *getPermAdjCOs_fixedDistr_2measures.R,* and *compareAdjDistanceCombine2Measures.R*)

We also calculated crossover interference in terms of each donor's individualized genetic map (**Extended Data Fig. 11e**). We determined the proportion of cells with a second crossover in windows of sizes 5–95 centimorgans on one chromosome at a time (containing 5–95% of the total crossovers from cells with two crossovers on that chromosome). Starting with each crossover on any chromosome with at least 30 crossovers observed across all cells, we identified the window containing X% of the rest of the crossovers on that chromosome in that individual. (If this crossover was near the end of the chromosome such that such a window was impossible, it was dropped from analysis, although it would have been included in previous crossovers' windows.) We noted whether this chromosome's second crossover fell in this window. We did this for each two-crossover chromosome (*n* per donor noted above), and then determined the proportion of cells with a second crossover in this cM window and compared it to the window size (*i.e.*, at a window size of 5 cM, or 5% of all crossovers from two-crossover chromosomes, far fewer than 5% of cells contain a second crossover). We then compared the observed percentage at each expected percentage (in each 5-cM window) across individuals, both visually and using the Kruskal–Wallis test (**Extended Data Fig. 11f**). To confirm that the results were not dependent on the direction of analysis or the specific crossovers in each window, we implemented this analysis going both from "left" to "right" (increasing physical position) and from "right" to "left" (decreasing physical position) on a chromosome. (See our script *computeSuppression.R*.)

High–crossover rate donors may have a different chromosomal composition of two-crossover chromosomes than low–crossover rate donors, *e.g.*, few two-crossover chromosome 1s but many two-crossover chromosome 18s, while low crossover rate donors may have the reverse. To determine whether the observation of individuals' crossover interference differences and the negative correlation of interference with crossover rate was robust with respect to this differential composition, we down-sampled each individual to have the same number of two-crossover chromosomes for each chromosome as the individual with the lowest number of two-crossover chromosomes of that number, and then repeated our analyses (*n* total two-crossover chromosomes = 5,522). (See our script *compareDonorsConsecDist_samenobschrs.R*.)

In theory, differences among individuals' crossover interference on chromosomes with two crossovers could be due to differential failure to detect crossovers at the very end of the chromosome. This would lead to the inclusion of chromosomes that actually had three crossovers, only two of which were detected, such that the included distance actually belonged to a three-crossover chromosome. These wrongfully included three-crossover chromosome distances would be shorter on average than two-crossover inter-crossover distances. Such mistaken inclusion of three-crossover chromosomes could occur preferentially in higher–crossover rate sperm donors, because higher–crossover rate individuals would be more likely to have a third crossover that could be missed. If so, it could give rise to the observed interference-rate relationship. To determine how this might manifest, we preferentially removed 10% of the chromosomes with the shortest inter-crossover distances from the individual with the highest crossover rate, left all chromosomes in for the individual with the lowest crossover rate, and removed percentages of shortest distances weighted by crossover rate from the intermediate crossover rate samples. The choice of 10% was overly conservative, as it is more than double the fraction of crossovers that we expect to be missed based on biased coverage near the telomeres: the estimate from crossovers in bead doublets that are discordant and near the end of chromosomes ranged from 0.2–4.0% across donors and was 2.1% globally (**Extended Data Fig. 2e**) (*n* two-crossover chromosomes retained per donor = NC1: 5,337, NC10: 6,120, NC11: 104,57, NC12: 11,107, NC13: 8,450, NC14: 7,344, NC15: 9,171, NC16: 9,214, NC17: 8,186, NC18: 8,831, NC2: 8,268, NC22: 9,166, NC25: 12,392, NC26: 5,300, NC27: 7,019, NC3: 7,084, NC4: 8,084, NC6: 7,466, NC8: 9,144, NC9: 10,359). We then repeated the crossover interference analysis with these unequally downsampled chromosome sets. (See our script *controlSimTelBias_MultSampInterference.R*; we used prop 0.1 and method *corate.percentile* in the parameter file for the described analysis.)

**Analysis of crossover interference and proportion of crossovers in distal zones across sperm cells**

To determine whether increased crossover interference was associated with lower crossover rate in sperm cells, we first assigned each cell (within a donor) to a decile based on its crossover number. We then compared the distance between all consecutive crossovers on each chromosome with two crossovers from each cell in the bottom decile (*i.e.*, the 10% of cells with the lowest crossover rate) to the same measurements from each cell in the top decile (the 10% of cells with the highest crossover rate).

To determine whether increased crossovers in the most telomeric zones of chromosomes was associated with lower crossover rate in sperm cells, we determined the proportion crossovers in two-crossover chromosomes that occurred in end zones for each cell (sum of crossovers occurring in end zones on two-crossover chromosomes / sum of all crossovers occurring on two-crossover chromosomes). We then compared these proportions across the top and bottom crossover-rate deciles determined as described above.

To increase power, we aggregated all cells across all donors by converting each measurement to percentiles within donors: crossover number per cell and each proportion of crossovers occurring in end zones was converted to a percentile for that sample. We then combined all cells, re-computed crossover-rate deciles based on these combined percentiles, and performed comparisons across these crossover-rate deciles. For crossover interference, we took the percentile of each inter-crossover distance for each chromosome separately (then combined across chromosomes) to control for differences in the composition of two-crossover chromosomes among donors. These distance percentiles were compared in the Mann–Whitney test across crossover number deciles, and the median for each cell was plotted to show each cell's aggregate phenotype. (**Fig. 4d,e, Extended Data Fig. 16**) (See our script *rateVOtherPtypesAcrossCellAggs.R*, which performs these and other analyses. We used 10 for the 6$^{th}$ argument ["Number of groups to split cells into based on CO rate for 'meta-cell' analyses"] for the analyses described here.)

**Identification of aneuploidy and chromosome arm-scale structural variants**

We used an approach based on sequence read depth to determine copy number in regions across the genome and identified chromosomes or chromosome arms with aberrant read depth to identify aneuploidy. As described above (see "Restricting to cell barcodes with coverage of the entire genome"), we used Genome STRiP (http://software.broadinstitute.org/software/genomestrip/)[24,25] to determine read depth across the genome in each sperm cell.

Before looking for aneuploidy events, we first removed bins that had outlying read depth across all cells, defined as those with $p < 0.05$ in a one-sided one-sample $t$-test (looking for increased read depth) against the expected mean read depth of 2# (defined below). To identify gains of autosomes, we then performed a one-sided one-sample $t$-test (expecting increased read depth in a gain) for each cell against expected read depth for a gain of one copy, 2#. For each cell, this analysis compared the distribution all bins' read depth across a region of interest to the gain expectation 2#, and flagged any cells whose read depth distributions were not significantly different ($p \geq 0.05$) We used the same approach to identify losses, comparing a cell's read depth distribution across bins to 0.1 and flagging any that were not significantly higher ($p \geq 0.05$).

The expected copy number – and thus read depth – for gains is 2, but the expected read depth for gains depends on the size of the chromosome, because the total increased number of reads in a library with a gain compared to the number that would be in that same library without a gain pulls read depth down globally by increasing the total number of expected reads, causing the denominator in each read depth bin (the expected number of reads in that bin, tied to total number of reads) to increase. To correct for this effect of a gain itself on global read depth, we defined the critical value at or above which gains were identified as a chromosome-specific value, slightly below 2: 2# = 2*(the proportion of the genome in base pairs coming from all chromosomes other than the tested one). We used 0.1 rather than 0 because the expected read depth for losses, as a small number of reads generally align to a lost chromosome (due to mis-alignment or possibly DNA from different sources being present in the droplet).

For non-acrocentric chromosomes, we performed gain and loss calling for the arms separately, as well as for the whole chromosome. Because amplification of more than two copies of a chromosome arm could result in the whole chromosome passing the *p*-value threshold, we required a whole-chromosome event to both pass the *p*-value threshold at the whole-chromosome level and to have rounded read depth of both arms ≥ 2 for a gain (or 0 for a loss). For the acrocentric chromosomes (13, 14, 14, 21, 22, Y), only the *q* arm was considered and any *q* arm gain or loss was considered to be a whole-chromosome event due to the difficulty of processing the *p* arm (unless investigated further, *e.g.,* **Fig. 6c**). The total number of copies gained was inferred from the overall read depth for any flagged chromosome (**Fig. 6, Extended Data Fig. 19**).

For the sex chromosomes, we followed a similar statistical framework, but did not call a flagged loss as an aneuploidy unless losses were flagged for both the X and the Y chromosomes. A gain was also called if both the X and Y chromosomes were present (*i.e.* not flagged as losses). (See our scripts *setupgsreaddepth.R, idaneus_initialttests.R, curateaneudata_clean.R, getautosomalaneumatrix.R*, and *getxykaryos_aneus.R* for aneuploidy calling and output formatting; see our scripts *curateAnFreqFromCodeMatrix.R, curateInitAnalyzeXYKaryos.R,* and *combineAnFreq_AutXY.R* for conversion of outputs of aneuploidy calling to cross-donor aneuploidy frequency tables.)

**Identification of the meiotic division of origin for chromosome gains**

To see whether chromosome gains originated in meiosis I (MI) or meiosis II (MII), we determined whether the centromeres of the multiple copies of the chromosomes were heterozygous and therefore from homologous chromosomes, which typically disjoin in MI, or homozygous and therefore from sister chromatids, which typically disjoin in MII. We first identified heterozygous regions for all cells using a Hidden Markov Model (HMM) in which the states are 1) heterozygous (emitting either haplotype's alleles) or 2) homozygous (emitting only one haplotype's alleles), with transition probability between the states equal to the recombination transition probability (see "Identification of crossover events" section), saving the start and end positions and indexes of any heterozygous tracts of SNPs. For each gain, we then determined whether heterozygous tracts overlapped with the centromere, with the same centromere locations as those used for SNP calling (from the UCSC cytoband track[19,20]). If a heterozygous tract 1) started before the start of the centromere and ended after the end of the centromere or 2) started at the first SNP observed on an acrocentric chromosome or within the first 10 SNPs and was more than 10 SNPs long, it was classified as an MI gain; if no heterozygous tract overlapped the centromere, it was classified as an MII gain. (See our scripts *getDiploidTracts_hmm.R, originOfGainID.R,* and *curateOriginMultSamps.R*.)

For the sex chromosomes, we used the logic that the X and Y chromosomes are homologs and separate at MI, whereas X and Y sisters separate at meiosis II. Therefore, any XY sex chromosome gain derives from MI, whereas an XX or YY gain derives from MII.

**Examination of the relationship between recombination and aneuploidy**

We examined the relationship between recombination and aneuploidy at three levels: sperm donor, cell, and aneuploid chromosome. To determine whether the aneuploid chromosomes themselves had fewer crossovers than chromosomes that were not lost or gained, we first determined the number of crossovers on chromosomes that had been gained by identifying the number of transitions between heterozygous and homozygous states using an

HMM, as described above in "Identifying the meiotic division of origin for chromosome gains." This is the total number of gains that occurred on both of the present chromosomes together, as it is impossible to determine for, *e.g.*, two crossovers, whether one occurred on each starting chromatid or both occurred on one starting chromatid. (See our script *getGainChrCOs.R*)

This process sometimes yielded very many crossovers (>10) being called on gained chromosomes because the presence of two haplotypes can be difficult to algorithmically distinguish from multiple crossovers depending on the haplotype patterns of observed alleles make. Therefore, we performed downstream analyses on 1) all gained chromosomes, including those with these high crossover numbers and 2) on the large majority of gained chromosomes with fewer than five called crossovers to exclude any with a crossover number that was likely to be inflated. We report the results of both versions of the analysis in the supplemental text, and the result of the analysis excluding chromosomes with inflated crossover number in the main text and figures.

We then calculated the total number of crossovers occurring on all gained chromosomes, chromosomes gained in MI, and chromosomes gained in MII; all donors' gains of one copy were included. To determine whether these numbers were lower or higher than expected, we ascertained 10,000 matched sets of the same number of gains and compared the sum of crossovers for each of the sets to our observed total (**Fig. 5f, Extended Data Fig. 18a**), computing a one-sided *p*-value based on the hypothesis that gained chromosomes would have fewer crossovers. For each matched gain, we considered each chromosome gain, randomly selected two non-aneuploid cells from the same donor, and summed the crossovers on the same chromosome as the gain, thereby controlling for differences in crossover rate among chromosomes and individuals. In each matched set, we performed this procedure for each of the observed gains and summed all crossovers. (See our script *combineGainsLookInCis.R*.)

To determine whether cells with aneuploidy had fewer crossovers overall on the remaining, non-aneuploid chromosomes than euploid cells, we first determined the number of crossovers per non-aneuploid megabase in each cell in order to control for aneuploid territory: for euploid cells, all chromosomes were included, whereas for aneuploid cells, aneuploid chromosomes were excluded. The set of euploid cells used for comparison against aneuploid cells included only cells with no detected structural variant, including arm-level chromosome gains or losses. In each sperm donor, we used the Mann–Whitney test to compare the distribution of crossovers per megabase in cells with any aneuploidy, MI gains, or MII gains to the distribution of crossovers per megabase in euploid cells. To increase power, we pooled all cells from all donors, taking the within-donor *z*-score of crossovers per megabase to control for crossover rate differences among donors, and repeated the same tests (**Extended Data Fig. 18b**). To demonstrate that we could detect differences between aneuploid and euploid cells without correcting for aneuploidy (when aneuploid chromosomes' 0 crossovers were included in the analysis), we performed this analysis on the total number of crossovers per megabase in the genome, rather than non-aneuploid territory. To assess in a different way whether aneuploid status alone, rather than the absence of the chromosome from analysis due to the aneuploidy, was significantly associated with crossover number, we also performed a linear regression including all cells as observations, using the following equation:

### Crossovers = [any whole chromosome aneuploidy: 0 = no; 1 = yes] + [0 of 1 for aneuploidy at each chromosome: 1 = aneuploidy] + [sperm donor dummy variables with values of 0 or 1 to control for underlying differences in crossover and aneuploidy frequency]

We performed this analysis without chromosome covariates to demonstrate that we did have power to detect a relationship at the level of entire chromosomes left out of aneuploid cells. (See our scripts *coPerMbVaneuploidy.R, linregCOVAneuploidy.R,* and *mImIIgains_copermbandlinreg.R*)

At the donor level, we performed a Pearson's correlation test of mean crossovers per cell per donor versus mean (whole-chromosome) aneuploidy events per cell per donor, the mean MI gains per cell per donor, and the mean MII gains per cell per donor (**Extended Data Fig. 18c**). We calculated the statistical power for this analysis using the function *pwr.r.test* from the R package *pwr* (https://CRAN.R-project.org/package=pwr).
